# Direct quantitative assessment using digital droplet PCR and field-scale spatial distribution of *Plasmopara viticola* oospores in vineyard soil

**DOI:** 10.1101/2024.07.29.605284

**Authors:** Charlotte Poeydebat, Eva Courchinoux, Isabelle D. Mazet, Marie Rodriguez, Alexandre Chataigner, Mélanie Lelièvre, Jean-Pascal Goutouly, Jean-Pierre Rossi, Marc Raynal, Laurent Delière, François Delmotte

**Affiliations:** Bordeaux Sciences Agro, INRAE, SAVE, ISVV, Gradignan, France; INRAE, Bordeaux Sciences Agro, SAVE, ISVV, Villenave d’Ornon, France; Université Bourgogne Franche-Comté, INRAE, AgroSup Dijon, Agroécologie, Dijon, France; INRAE, Université de Bordeaux, Bordeaux Sciences Agro, EGFV, ISVV, Villenave d’Ornon, France; INRAE, CIRAD, IRD, Institut Agro, CBGP, Montferrier-sur-Lez, France; IFV, Bordeaux Nouvelle Aquitaine, UMT SEVEN, Villenave d’Ornon, France

**Keywords:** oomycete, downy mildew, primary inoculum, geostatistics, ddPCR, bioassay, soil infectious potential

## Abstract

Grapevine downy mildew, caused by the oomycete *Plasmopara viticola*, is one of the most devastating diseases of grapevine worldwide. Primary inoculum (*i.e.* oospores) play a decisive role in downy mildew epidemics, but we still know very little about its abundance in vineyard soil. This study presents a novel molecular method for quantifying *P. viticola* oospore concentration in vineyard soil using digital droplet PCR (ddPCR). The development of this method enabled characterization of both the abundance and spatial distribution of oospores in a vineyard at the onset of the growing season. Following a regular grid, a total of 198 soil samples (0-15cm horizon) were collected in March 2022 in grapevine rows in a 0.22 ha vineyard planted with cv. Merlot and conducted according to French organic viticulture specifications. Additional samples were collected from the same field within five nested sampling plots with three distance levels, including samples collected in the inter-rows. Using ddPCR, we found *P. viticola* DNA in all soil samples except one, and we estimated that oospore concentration ranged from 0 to 1858 oospores per gram of soil (303 ± 308 on average). The distribution of oospores at field scale was not random but characterized by 15 m-diameter patches of concentrically increasing oospore concentration. Oospores accumulated 5 times more below the vine stocks than in the inter-row. Using a leaf disc bioassay, we found that soil infectious potential significantly increased with oospore concentration assessed by ddPCR. However, the low coefficient of determination of the relationship indicated that DNA-based oospore quantification lacked clear epidemiological significance. Both ddPCR and bioassay methods are valuable tools that could be used to assess reservoirs of *P. viticola* primary inoculum across different agroclimatic contexts, thereby bringing greater genericity. Further methodological improvement will also help refine the accuracy of DNA-based assessment of primary inoculum reservoir and improve our understanding of the relationship between primary inoculum reservoir and epidemic dynamics. Ultimately, these data will be essential for improving epidemic risk models and evaluating new preventive disease management strategies targeting the primary inoculum.

**Importance:** Grapevine downy mildew caused by the oomycete *Plasmopara viticola*, affects leaves and bunches, and leads to important economic losses for viticulturists. Recently, evidences have accumulated that soilborne primary inoculum (*i.e.* oospores in the soil) importantly contributes to disease progress. The significance of our work is in presenting a direct and sensitive method for assessing soil oospore concentration, as well as quantitative and spatially-explicit data on downy mildew primary inoculum. This opens the way to new research, the evaluation of new disease control strategies based on primary inoculum management and the improvement of epidemic risk models, which will potentially contribute to lower fungicide use in viticulture *in fine*.

## Introduction

While crop protection from pest and diseases is largely ensured by the use of pesticides worldwide (Sharma et al., 2019), there is accumulating evidence that they are responsible for environmental pollution threatening both biodiversity (Wan et al., 2025; Beaumelle *et al*., 2023; Beketov *et al*., 2013) and human health (*e.g.* Dall’Agnol *et al*., 2021). In Europe and France, policies are clearly encouraging the reduction of the use of chemical plant protection products (European Green Deal and National Action Plan Ecophyto; Lamichhane *et al*., 2019), either by improving use efficiency or by developing alternative pest and disease management strategies. Grapevine downy mildew, caused by the obligate biotrophic oomycete *Plasmopara viticola*, is one of the most devastating diseases of grapevine worldwide. Its management still relies mainly on intensive use of fungicides throughout the growing season. In 2006, the French wine-growing area covered 3.3% of utilized agricultural land but contributed up to 14.4% of national fungicide use (Butault et al., 2011). In 2019, more than 80% of the treatment frequency index was made up of fungicides used against downy mildew and powdery mildew (Agreste, 2023).

In temperate regions, the life cycle of *P. viticola* comprises a phase of asexual multiplication on its host tissues during the growing season and a phase of sexual reproduction, which occurs in infected grapevine leaves between thalli of sexually compatible strains (as *P. viticola* is heterothallic, Wong *et al*., 2001) leading to the formation of oospores. Leaves then fall on the ground and oospores overwinter in leaf debris or as free propagules in the soil. The stock of oospores in the soil constitutes the primary inoculum for the next growing season, which will be responsible for the onset of the epidemics. After the first primary contaminations have occurred, epidemic development depends on both primary and secondary infections, whose relative contributions depend on climatic conditions of the season (*e.g.* Rumbou and Gessler, 2004). However, primary inoculum is increasingly considered to play a more important role in grapevine downy mildew epidemics than previously thought (Gessler *et al*., 2011; Gobbin *et al*., 2005) for several reasons. First, population genetic studies evidenced that grapevine downy mildew populations are panmictic across European vineyards (Fontaine *et al*., 2021, 2013) showing that genetic mixing resulting from sexual reproduction largely structures pathogen populations. Second, in contrast to the widely-held assumption that primary contaminations rarely occur after flowering, it has been shown that oospores are able to germinate and cause infections throughout the growing season (until late September in the northern hemisphere; Kennelly *et al*., 2007; Maddalena *et al*., 2021) and that a substantial amount of infections, including in the middle of the growing season, can arise from oospores (up to 40% of contaminations in the most favorable conditions; Gobbin *et al*., 2003; Hong *et al*., 2020). Third, while the epidemic continues to progress after harvest (from what we observe in surveyed vineyards), the incidence of typical mosaic-like downy mildew symptoms on grapevine leaves was shown to be proportional to the amount of oospores formed in the leaves (Si Ammour *et al*., 2020). Finally, downy mildew incidence at the end of the season was found to explain the precocity and intensity of the disease the following season in Canadian vineyards (Carisse, 2016).

While actual control of grapevine downy mildew primarily targets the asexual propagation of the disease, the contribution of *P. viticola* primary inoculum to epidemic dynamics suggests that measures targeting the primary inoculum (*e.g*. post-harvest leaf removal, biosolutions preventing sexual reproduction or oospore survival, practices limiting oospore germination or splashing) could contribute to downy mildew management, in particular under circumstances favorable to primary contaminations. At least, decisions on the application of fungicides should better account for the abundance of primary inoculum and its contribution to epidemic risk. Although the stages of the sexual reproduction cycle of *P. viticola* are well known (see for example Gessler *et al*., 2011), their quantitative description remains incomplete. The study of disease inoculum abundance and spatial distribution in agricultural soils are generally restricted to soilborne pathogens, including oomycetes (see for example Botero-Ramirez *et al*., 2021; Moussart *et al*., 2009). To our knowledge, these questions have rarely been addressed regarding strictly airborne biotrophic pathogens with a telluric conservation life stage. Molecular approaches have been developed and used to quantify airborne asexual inoculum of grapevine downy mildew, as well as oospores in grapevine leaves (Carisse *et al*., 2021; Douillet *et al*., 2022; Si Ammour *et al*., 2020; Valsesia *et al*., 2005). Direct indicators of the pathogen abundance in the vineyards, such as spore concentration in the air, are progressively included in decision tools or regional monitoring programs (Brischetto *et al*., 2020; Douillet, 2023). Yet, the prediction of downy mildew epidemic risk in decision support systems is mainly based on meteorological variables that control the development of *P. viticola* (Dubuis *et al*., 2012; Park *et al*., 1997; Rossi *et al*., 2008a; Strizyk *et al*., 1983). Indicators of primary inoculum reservoir in the soil compartment have never been accounted for until now, partly because quantitative data is scarce. Instead, predictive models assume a fictitious quantity of primary inoculum and calculates rates of change (associated with maturation, dispersal or germination; *e.g.* Rossi *et al*., 2008a). Recently, a molecular method based on real-time qPCR was used to quantify oospores in soil of vineyards in China (Yang *et al*., 2023). Quantitative data on oospores obtained from such methods support the transition to more sustainable downy mildew management in viticulture by improving epidemiological models for epidemic risk prediction, and enabling evaluation of control strategies based on primary inoculum management.

Here, we present a direct and sensitive method for the quantification of *P.viticola* oospores in vineyard soil based on the quantification of *P. viticola* DNA by digital droplet polymerase chain reaction (ddPCR). We provide an estimation of the number of copy of the ITS sequence used for amplification (ITS1/5.8S) in the genome of *Plasmopara viticola* that we used to convert the ddPCR output into a number of oospores per gram of soil. By means of geostatistics, we described the spatial distribution of downy mildew primary inoculum in a vineyard at the onset of the growing season. Finally, we build on previous works to conduct a bioassay to assess the infectious potential of soil samples and discuss the epidemiological significance of DNA-based assessments of downy mildew primary inoculum.

## Material & Methods

### Soil sampling and preparation

We conducted our study in an experimental vineyard with a sandy-gravel soil located at the INRAE Bordeaux Nouvelle-Aquitaine research center (Villenave d’Ornon, France; 44.79194, −0.57702) within the Pessac-Léognan appellation area. The surveyed vineyard was planted in 2011 with 18 rows of 68 Merlot grapevine stocks over a 0.22 ha surface (70 x 32 m), corresponding to a density of 6600 vines per ha (*i.e*., 1.6 m between rows and 0.95m between vines along the row). It was managed according to French organic farming specifications, meaning that Bordeaux mixture (copper) was the only chemical fungicide allowed to control downy mildew. Inter-rows were either sown with a mix of plant species selected for green manuring or covered by spontaneous vegetation in an alternating arrangement rotating every year. All inter-rows were tilled before grapevine budburst. For winter protection and weed management purposes, soil from both sides of the row was turned over on the lower part of the vines in the fall. The ridge formed under the row was then reopened in the spring. The surveyed vineyard was bordered on each of its long sides by a diversified shrubby hedgerow (*ca*. 1.5-2 m high) that separated it from other experimental vineyards.

Soil sampling was done in March 2022. We collected 198 soil samples at the nodes of a 2.85 x 3.2 m regular grid covering the entire surface of the vineyard; *i.e.* one sample every three vine stocks in the row, in one row out of two (Figure 1). To explore the spatial structure of primary inoculum of downy mildew in the soil at a finer scale, we delineated five plots in the experimental vineyard within which we collected soil samples following a nested sampling design with three distance levels. Each plot consisted of a 2.85 x 3.2 m rectangle containing 28 sampling points (Figure 1). The first distance level corresponded to the four corners of the plot (n_3_=4), hence sampling points were separated by about 3 m (samples were the same as for the regular grid sampling). The second distance level was set at 0.8 m (half the inter-row width), and sampling points were located in the corners of four 0.8 m squares located in each corner of the plot (n_0.8_=16, including 4 points in common with the first distance level; see Figure 1). The third distance level was set at 0.2 m with sampling points corresponding to the corners of four 0.2 m squares located in the corners of one of the four 0.8 m squares (n_0.2_=16, including 1 point in common with the first distance level and 4 points in common with the second distance level; see Figure 1). We also considered that the points from the regular grid were located in the row, while the points from the 2^nd^ and 3^rd^ levels of the nested sampling were located in the inter-row space. At each sampling point of both the regular grid and the nested sampling plots, we collected 500 g of soil from the top 15 cm using an auger. We ended with 318 soil samples, including 198 samples from the regular grid and 120 additional samples from the five nested sampling plots.

**Figure 1.**
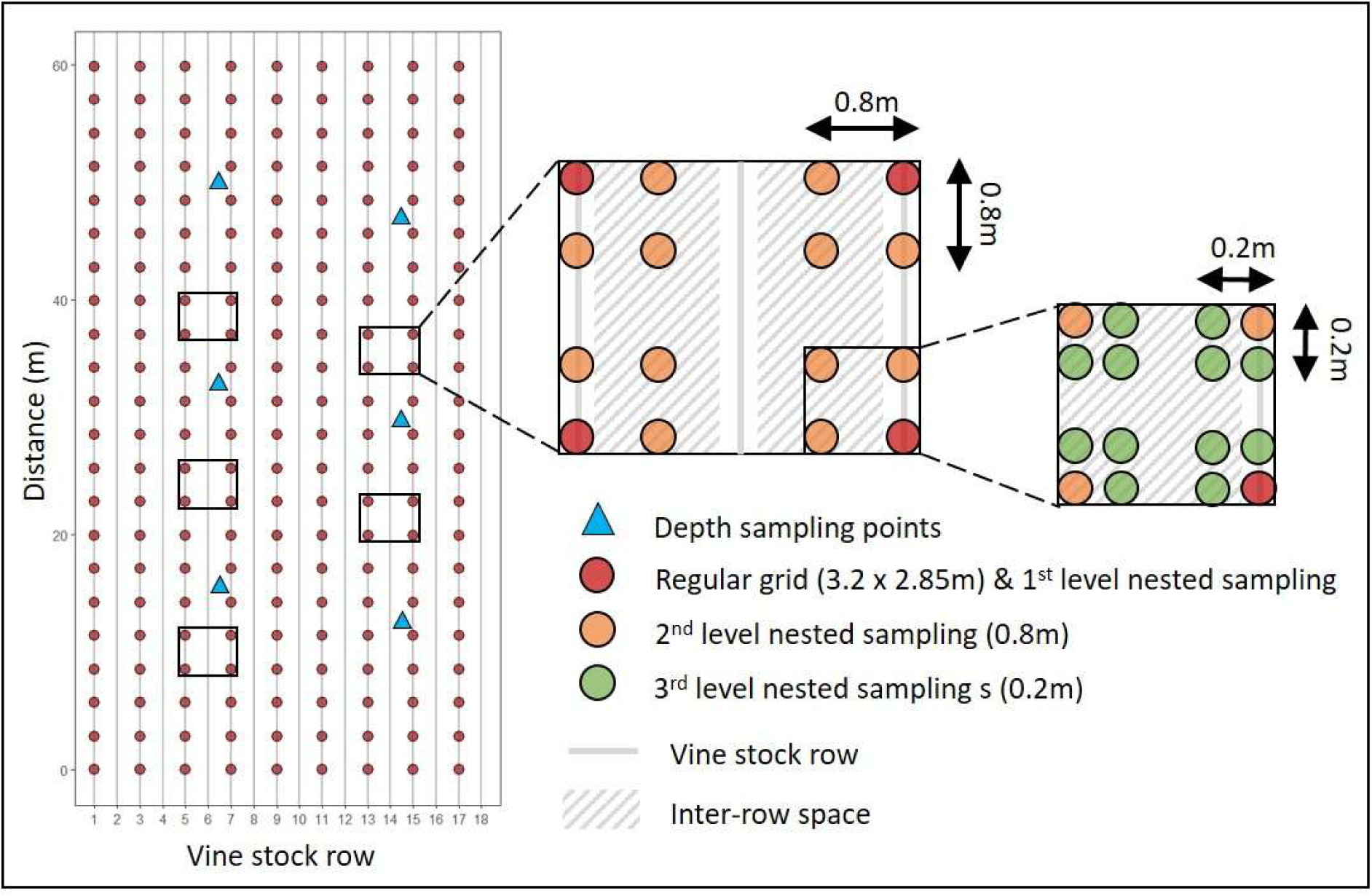
Map of the soil sampling points within the experimental vineyard.

To investigate the vertical distribution of *P. viticola* primary inoculum in the soil, we dug a hole 40 cm deep and 30 cm in diameter at six points located in the inter-row space in the experimental vineyard (Figure 1). We divided the soil profile into four strata of 10 cm (0-10 excluding leaf debris on soil surface, 11-20, 21-30 and 31-40 cm) and sampled 500 g of soil from each stratum, starting at the bottom of the profile to avoid contaminating deeper layers with upper ones. This corresponded to 24 samples in total (4 samples per hole or sampling location). All soil samples were stored at 5°C before processing.

### Quantification of *Plasmopara viticola* primary inoculum in vineyard soil samples

#### DNA extraction

Each sample was passed through a 4 mm mesh sieve and thoroughly homogenized for 30 s using an electric mixer set at medium speed. Directly after homogenization, *ca*. 6 g of the sample was collected and placed in a 5 ml tube to be freeze-dried. Then, a 2 g subsample of dry soil was placed in a 15 ml Falcon tube containing 4 g of 0.1 mm diameter silica beads, 5 g of 1.4 mm diameter ceramic beads and eight 4 mm diameter glass beads (as recommended by MP-Biomedicals for TeenPrep™ Lysing Matrix E). For the grinding and lysis extraction step, a volume of 8mL of lysis buffer composed of 100 mM Tris-HCl (pH 8), 100 mM EDTA (pH 8), 100 mM NaCl, 20% SDS and ultra-pure water was added in each tube. Mechanical cell lysis was achieved placing the tubes in a FastPrep-24 classic instrument (MP-Biomedicals) set for three runs of 30 s at 4 m.sec^-1^. Tubes were then incubated for 30 min at 70°C to activate chemical lysis and centrifuged at 5000 g for 7 min at 20°C to eliminate particles in suspension. DNA extraction was carried out using 1 ml of supernatant obtained from the centrifugation, as follows: 1) *Deproteinization:* addition of 100 µl potassium acetate 3M pH5.5 to the sample, homogenization and incubation on ice for 15 min to precipitate proteins, centrifugation at 14000 g for 5 min at 4°C to eliminate precipitated proteins (supernatant recovery and pellet disposal); 2) *DNA precipitation:* addition of 900µl cold isopropanol (v/v) to the supernatant from the deproteinization, gentle agitation of the tubes that were then placed at −20°C for 30 min, centrifugation at 13000 rpm for 30 min at 4°C, elimination of the supernatant; 3) *DNA Washing:* rinsing of the pellet with 400µl 70° cold ethanol, centrifugation at 13000 rpm for 5 min at 4°C, elimination of the ethanol supernatant, drying of the pellet by placing the tubes in a drying oven at 60°C for 20 min; 4) *DNA Purification:* resuspension of the pellet in 500 µl of ultra-pure water. Raw DNA extracts were further purified using a NucleoSpin Soil kit (Macherey-Nagel) following the manufacturer’s protocol from step 4 (addition of SL3 buffer). Finally, purified DNA extracts were eluted in 60 μL of buffer (TrisHCL, 5mM, pH 8.5) and stored at −20°C.

#### Digital droplet polymerase chain reaction (ddPCR)

To avoid common PCR inhibition problems with the soil matrix and to obtain direct absolute DNA quantification (no standard curve needed), samples were analyzed by droplet digital PCR (ddPCR) using the QX200 ddPCR system (Bio-Rad) composed of a droplet generator, a thermal cycler and a droplet reader. We used the specific PCR primers and TaqMan probe (Giop set) designed by Valsesia *et al*. (2005) that target the ITS1/5.8S to amplify *P. viticola* DNA in our samples.

For each sample, we added 4 µl of DNA extract diluted to the tenth to 18 µl of a PCR mix composed of 11 µL of 2x ddPCR Supermix for Probes (Bio-Rad), 2.2 µL of 7.5 µM Giop F, 2.2 µL of 7.5 µM Giop R, 2.2 µL of 5 µM Giop P-VIC and 0.4 µL of ultra-pure water. The samples (20 µl) were first placed in the droplet generator to proceed to the breakdown of the samples into 15000 to 20000 micro-droplets using oil. Samples were then carefully transferred to 96-well plates with a pipette, with one sample per well (one single analysis per sample). The sample plates were placed in the thermal cycler for DNA to be amplified according to the following program: 1) an initial phase at 95°C for 5 min; 2) 40 cycles of 30 s at 95°C then 1 min at 60°C then 30 s at 72°C; and 3) a final phase at 98°C for 10 min. Samples were cooled at 12°C before being transferred to the droplet reader for DNA quantification (ratio of positive *vs* negative droplets). Finally, we obtained one value per sample that corresponded to the number of ITS1 copies per µl of reaction mix.

#### Performance of the method to quantify P. viticola oospore DNA in soil

In order to determine the performance of the overall method to quantify *P. viticola* oospore DNA in soil, we spiked and analyzed non-infected 2g soil samples with *P. viticola* oospores at different known concentrations (see details in Supplementary Materials, Box S1 and Table S1). In particular, we assessed the linearity (using a linear regression analysis) and the efficiency (as the percentage of expected DNA actually measured) of the method. In addition, we conducted a separate analysis on two subsets of the soil samples taken from the vineyard to assess the repeatability of the DNA extraction and ddPCR assay steps on environmental samples naturally containing oospores (see Box S2 in Supplementary Materials for details).

#### Estimation of ITS1/5.8S copy number in Plasmopara viticola genome

In order to estimate the ITS1/5.8S copy number in the genome of *P. viticola*, we compared the ddPCR signals obtained with the ITS primer pairs (Valsesia *et al*., 2005) *vs* primers targeting the single copy β-tubulin gene (Weerakoon *et al*., 1998). First, sporangia of seven *P. viticola* strains frequently used in laboratory for genetic applications were collected with a paintbrush from grapevine (*Vitis vinifera* cv. Cabernet Sauvignon) leaves infected beforehand on purpose and placed into separate 2 mL microcentrifuge tubes with 1 mL Tris-EDTA buffer. Second, freeze-dried samples were extracted following a SDS based method (adapted from Möller *et al*., 1992) and DNA quantity and quality were checked by dosage with either Qubit 4 or Denovix-11. About 35 sequences from NCBI (Fontaine *et al*., 2021; Rouxel *et al*., 2013) were aligned with *P. viticola* genome (from Paineau *et al*., 2024) using Geneious Prime. Since none existed yet, β-tubulin primers (Forward: ACAGCCAATTTTCGTAAGTCC, Reverse: GCCTTGTACGACATTTGCTTT) and a TaqMan probe (5’ FAM-TGTCCGGGAAATCGAAGGCA-MGB 3’) were designed using the consensus sequence with 100% similarity. Specificity was tested using NCBI Blast (blast.ncbi.nlm.nih.gov) and by running quantitative PCRs on either pure *P. viticola* strain, *P. viticola* with grapevine leaf or grapevine leaf only (see Figure S2 in Supplementary Materials) using a QuantStudio 5 (Life Technologies). Reaction mix was composed of 10 µL of 2X PCR Buffer (Takyon Low Rox Probe MasterMix - Eurogentec), 1,6 µL of each β-tubulin primer (forward and reverse) at 10 µM, 0,4 µL of probe at 10 µM, 3,9 µL of molecular biology water (Sigma) and 2,5 µL of DNA template for a 20 µL final volume. Due to a large copy number difference between both genes and ddPCR sensitivity, we run the reactions for each marker in separate reaction wells. In consequence, each sample were normalized with a final quantity of 5 pg for ITS samples and 5 ng for β-tubulin samples. All reactions were run with a cycling program starting with 3 min at 95°C followed by 40 cycles of 10 s at 94°C alternating with 60 s at 60°C. For each strain, the number of ITS1/5.8S copies was estimated as the ratio of β-tubulin to ITS copy number per ng.

### Spatial structure of *Plasmopara viticola* primary inoculum in a vineyard

The spatial structure of *P. viticola* primary inoculum in the experimental vineyard was investigated at both field-scale and plot-scale by analyzing the regular grid data and the nested sampling plots data, respectively, using a geostatistical approach. Hereafter, we describe the field-scale analysis only, but the same methodology (except model fitting and kriging) was applied for the plot-scale analysis (see Table S3 in Supplementary Materials for details).

*Plasmopara viticola* DNA in soil samples, expressed as the number of ITS copies per µl of reaction mix, was log-transformed before analysis to reduce the skewness of the distribution and limit variance inflation and variogram distortion (Oliver and Webster, 2014). First, all pairs of data (N=19503) were segregated into distance classes. The maximal distance between two points in our dataset was 65.09 m. In our analysis, however, we considered a maximal separation distance of 60 m to exclude highest distance classes with a number of point pairs below the empirical threshold for semivariance estimates accuracy (at least 30 to 50 pairs required; Chiles and Delfiner, 1999; Journel and Huijbregts, 1976). We considered a spatial lag of 2 m, which led to a total of 29 distance classes, with 79 to 1498 pairs each. The sampling points in the first distance class were separated by a median distance of 3.02 m. We then computed the semivariance for each distance class using the Matheron estimator (Matheron, 1963), defined as follows:

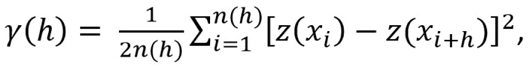

where *z*(*xi*) is the primary inoculum at the first point of the ith pair, *z*(*xi* + *h*) is the primary inoculum at the second point of the ith pair, h is the median of the class of distance considered and n is the number of point pairs in the distance class considered.

We tested for anisotropy in the data (*i.e.* spatial variation depending on direction) by computing directional experimental semivariograms (0°, 45°, 90° and 135°). Results revealed no clear anisotropy (see Supplementary Materials Figure S3 and Table S4). Hence, we represented the omnidirectional experimental semivariogram of the primary inoculum of *P. viticola* in the vineyard by plotting the semivariance against the sampling point separation distance h. A semivariogram shows how the similarity between samples changes with increasing separating distance *i.e.* spatial dependence. It can take various forms, and semi-variance often increases with distance until it reaches a plateau at which point it remains stable. The variogram is flat in the absence of spatial structure.

To determine the theoretical model best fitting our experimental semivariogram, we used the ‘autofitVariogram()’ function of the ‘autofit’ package in R to fit different variogram models to the data and calculate the sum of squared errors (sserr) between the experimental semivariogram and each fitted model (see Supplementary Materials Table S4). Using the sserr as the criterion to select the best-fitting model (Hiemstra *et al*., 2009), we choose to fit a Matérn model (with a smoothness parameter equals to 0.2) to the experimental semivariogram using ordinary least square residuals method (Cressie, 1985). The Matérn model *F*(*h*) and the associated semivariance *γ*(*h*) are as follows:

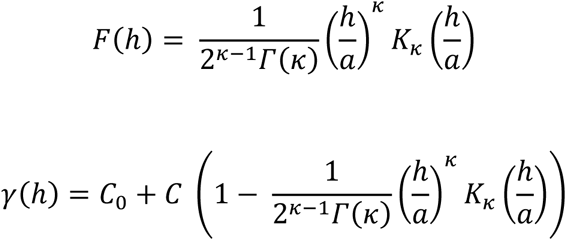

where *h* is the separation distance, *K_κ_* is a modified Bessel function of the second kind of order *κ* (Abramowitz and Stegun, 1972), *Γ* is the gamma function, and *κ* is the smoothness parameter. The model parameter estimates were regarded as quantitative descriptors of the spatial pattern of the data. They included i) the spatial range *a*, or the distance beyond which variance is constant; ii) the nugget variance *C*_0_, an estimate of error measurement plus error due to uncontrolled variability at distance lags shorter than the shortest lag considered; iii) the structural variance *C*; and iv) the sill *C*_0_ + *C*, or sample variance.

Before proceeding to kriging, we checked for the possibility of a trend in the data by comparing our raw experimental semivariogram to both a linear trend model (x + y) and a second order polynomial (quadratic) trend model (x + y + x^2^ + y^2^). The linear trend had a negligible effect on the empirical semivariogram (see Supplementary Materials Figure S4), indicating the absence of a significant large-scale drift. In contrast, the quadratic trend model effectively removed the spatial structure, resulting in an almost flat variogram. This suggested that the quadratic trend absorbed not just a trend but the spatial variability itself. Removing it would therefore eliminate meaningful spatial information. Based on these results, we concluded that explicit trend removal was not appropriate for our dataset. Consequently, we applied ordinary kriging to preserve spatial dependence and ensure reliable spatial predictions. We therefore used the shape and parameter estimates of the raw omnidirectional semivariogram model to interpolate *P. viticola* primary inoculum values at unsampled locations over a 0.05 m × 0.05 m grid by ordinary kriging, which is an optimal interpolation procedure described in detail in various sources (*e.g*., Goovaerts, 1998; Oliver and Webster, 2015). After back-transforming the resultant values, we produced a contour map of the spatial distribution of the primary inoculum in the field, as interpolated from our data (*i.e.*, given the local conditions and our sampling plan).

All geostatistical analyses were performed using the package gstat (Pebesma, 2004) in the R statistical software (v4.3.1; R Core Team, 2023). The contour map was designed using the ggplot2 package (Wickham, 2016).

### *Plasmopara viticola* primary inoculum along depth and distance-to-row gradients

In the nested sampling plots, the sampling points were located either below the vine stocks in the row (10 samples per plot) or in the inter-row space at 20, 60 or 80 cm from the nearest row (respectively 4, 4 and 10 samples per plot). We used this data to test whether the amount of primary inoculum in the soil varied along a gradient of distance to the row. We fitted a linear mixed model with the log-transformed number of ITS copies per µl as a response variable, the distance to the row as a fixed effect and the plot as a random intercept effect (to account for the dependence of the data within one plot). Model parameters were estimated by restricted likelihood estimation and the significance (α = 0.05) of the regression coefficients was tested with Student t tests and Satterthwaite’s approximation for degrees of freedom, using the lmer function of the lmerTest package (Kuznetsova *et al*., 2017). We evaluated model fit by calculating the percentage of variance explained by fixed (R^2^m) and by fixed plus random effects (R^2^c) (Nakagawa and Schielzeth, 2013). Finally, we tested whether the amount of primary inoculum varied with the depth of sampling using a Kruskal-wallis test and a Dunn test of multiple comparison. Statistical analyses were conducted using the R statistical software (v4.3.1; R Core Team, 2023).

### A bioassay to assess soil infectious potential

The principle of the bioassay is to place soil samples in optimal temperature and humidity conditions for oospore germination, to trap the released zoospores using susceptible grapevine leaf discs, and finally to approximate the infectious potential by evaluating the extent to which leaf discs are infected. The method described below was adapted from previous works (Hill, 1998; Pertot and Zulini, 2003; Rossi *et al*., 2008b).

We performed the bioassay on one eighth of our 318 soil samples (n=40) using a 1 g portion per sample. Each sample was placed in the bottom of a 10 cm diameter circular plastic box half-filled with osmosed water and incubated at 20°C in the dark to stimulate oospore germination (day 0). The sample was held at the bottom of the box using a piece of blotting cloth (80µm mesh). We collected leaves from untreated *Vitis vinifera* cv. Cabernet Sauvignon grafted plants (SO4 rootstock) grown in greenhouses. We specifically picked the third or fourth leaves from the apex, as leaves at this stage of development are particularly susceptible to downy mildew infection (Isabelle Demeaux, personal communication). We cut 10 mm diameter discs in the leaf blade with a manual round cutter. On the first day of the incubation (day 0), we placed 10 discs from 10 different leaves (to avoid biases due to differences in downy mildew susceptibility between leaves) on the surface of the water (abaxial face down) in each box. The leaf discs were used to trap the zoospores released into the water following germination of oospores present in the soil sample, *i.e.* the zoospores swam to the leaf discs and infected them via the stomata. The blotting cloth allowed the zoospores to pass through but prevented the discs coming into direct contact with the soil and limited contamination of the discs by saprophytic microorganisms. A separation structure was used to prevent the discs in a box from overlapping and maintain the same trapping surface in each box (see Figure S5 in Supplementary Materials). After 48 h, we placed the 10 discs abaxial face up on a moist Whatman® filter paper sheet in a transparent box sealed with plastic film (to maintain high humidity) and incubated at 20°C with a 12h-12h day-night light cycle for 6 days. After 6 days of incubation, we visually inspected the discs and evaluated the percentage of the area covered with *P. viticola* sporangiophores for each disc. The germination of a cohort of oospores under optimal conditions (immersed in water, in the dark and at 20°C) extend over 6 to 8 days, with a peak around the second and third days of incubation (Isabelle Demeaux, personal communication). To account for germination dynamics, we repeated zoospore trapping using leaf discs on day 2, 4 and 6 from incubation start (day 0). We used the cumulative number of infected discs and the mean percentage of leaf area infected as indicators of the infectious potential of the soil.

We assessed the relationship between the abundance of primary inoculum in the soil (as estimated with the ddPCR approach) and soil infectious potential by fitting one generalized linear model per indicator (response variable following Poisson distribution in both cases) with the number of *P. viticola* ITS copies per µl as a predictor. In both cases, model parameters were estimated by maximum likelihood and the significance (α = 0.05) of the regression coefficients was tested with z tests. We computed the coefficient of determination (R²) of the two models to determine the indicator best fitting the primary inoculum data. Statistical analyses were conducted using the R statistical software (v4.3.1; R Core Team, 2023).

## Results

### Quantification of *Plasmopara viticola* primary inoculum in vineyard soil samples

Considering the regular grid and nested sampling plot data, we found *P. viticola* DNA in all samples except one, with 6.62 ± 6.68 ITS copies per µl on average. The value for positive samples varied from 0.05 to 40.4 ITS copies per µl. A map of the raw data is available in Supplementary materials (Figure S6).

By comparing the ITS1/5.8S to the β-tubulin signal obtained by ddPCR for seven *P. viticola* strains, we found that the diploid genome of *P. viticola* contains on average 215 ± 37 copies of the ITS sequence, but varied from 175 to 281 copies depending on the strain (see Table S2 in Supplementary Materials for details). Using the average estimate for the number of ITS1 copies, we estimated that the number of oospores per gram of soil ranged from 0 to 1858 and that there was on average 303 ± 308 oospores per gram of soil in the experimental vineyard (see Figure S7 in Supplementary materials for computation details). Using the minimal and maximal estimates for the number of ITS1 copy, the average number of oospores per gram of soil was 372 ± 378 or 232 ± 235, respectively.

#### Performance of the method to quantify P. viticola oospore DNA in soil

*Plasmopara viticola* was detected at very low concentrations (*i.e.* 1 oospore in 2 g of soil; see Supplementary Materials Table S1) when analyzing spiked soil samples with our method. Different definitions of detection limit exist in the field of quantitative PCR (including ddPCR; see Hunter *et al*. 2017 for example), of which many correspond to a more or less conservative threshold rate of positive replicates (or detection frequency). In our study, *P. viticola* DNA was detected in 1 out of 5 replicates for the lowest concentrations (*i.e.* 1 and 10 oospores in 2 g of soil), and in 5 out of 5 replicates for higher concentrations (*i.e.* 50, 100, 500 and 1000 oospores in 2g of soil; see Supplementary Materials Table S1). This means that the detection limit of our method lies between 1 and 50 oospores per 2 g of soil, depending on the degree of conservatism applied. Besides, the analysis of the relationship between oospore concentration (or expected ITS copy number) and the ddPCR-assessed ITS copy number revealed that the method showed a good degree of linearity for the dynamic range tested in the present study (Pearson’s r = 0.97; see Supplementary Materials Figure S1). However, the degree of fit between the data and the linear regression was rather low for this type of analysis (R² = 0.93) and the number of ITS copies tended to be underestimated compared to what was expected (slope = 0.77). In addition, the efficiency of the method varied from 11 to 92% of expected DNA across the oospore concentrations tested (see Supplementary Materials Table S1). Consistently, the repeatability (expressed as within-sample coefficient of variation) of the DNA extraction and the ddPCR assay were found to be variable across vineyard soil samples (see Box S2 in Supplementary Materials for detailed results of the analysis of variance). On the positive side, median values for within-sample coefficients of variation were relatively low overall, for both the DNA extraction and the ddPCR assay (30% and 17%, respectively). Moreover, the analysis of variance also indicated that the error of measurement associated with the DNA extraction procedure and with the ddPCR assay were low compared to the between-sample variability. Altogether, these results indicate that although the repeatability and the efficiency of the method could be improved, it is useful to quantify P. viticola oospore DNA in vineyard soil.

### Spatial structure of *Plasmopara viticola* primary inoculum in a vineyard

After log-transformation, the skewness of the distribution of *P. viticola* DNA data was reduced from 1.69 to −0.35 for the regular grid, and from 2.40 to 0.42 for the nested sampling plots. The shape of the fitted Matérn model (regular grid data) revealed that the downy mildew primary inoculum was spatially structured at the field scale (Figure 2). The range was 15.83m. The nugget variance, the structural variance and the sill were 0.156, 0.354 and 0.510, respectively. The map of the values interpolated by ordinary kriging using the fitted model is shown in Figure 2.

**Figure 2.**
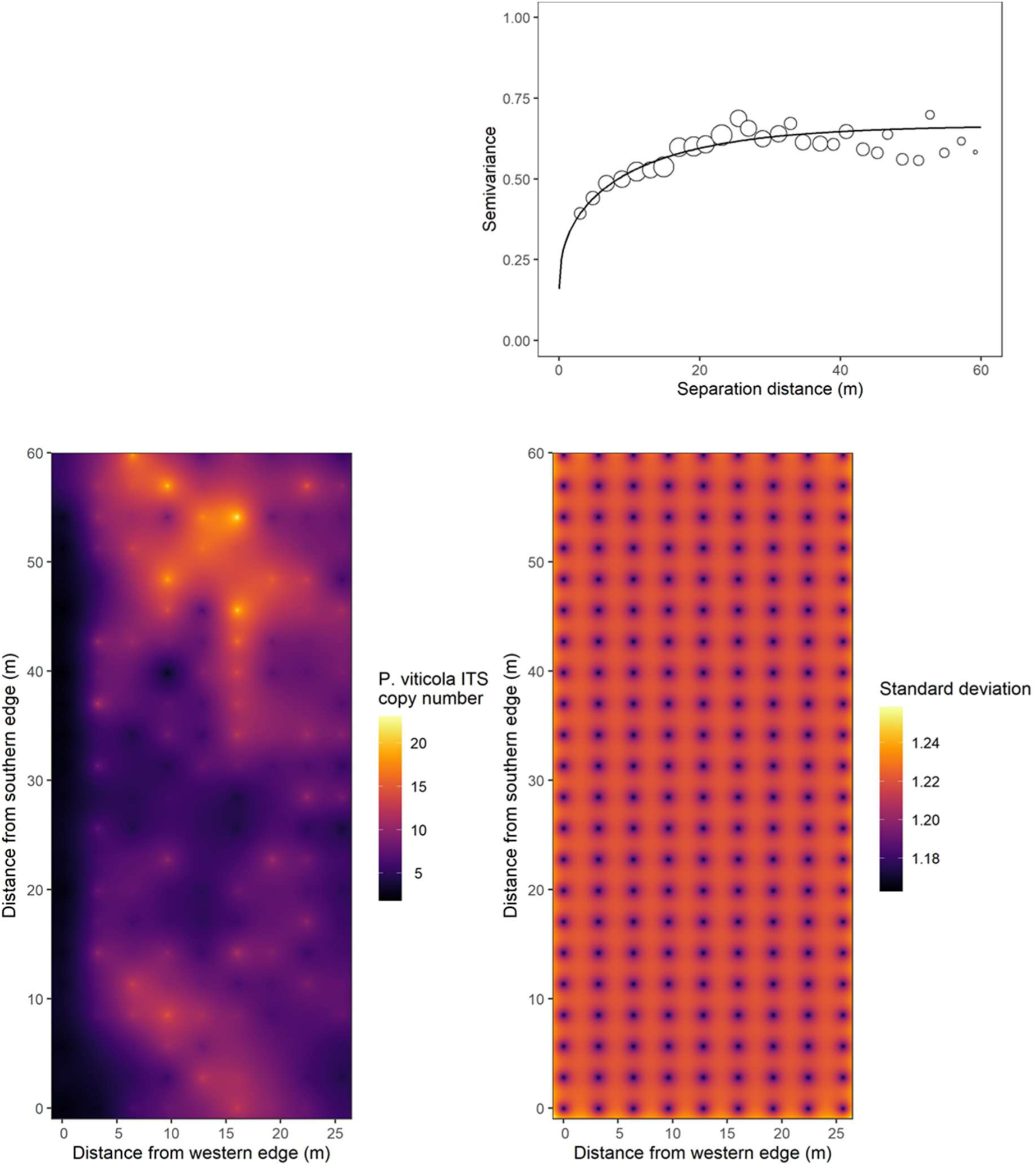
Experimental omnidirectional semivariogram of *Plasmopara viticola* DNA (expressed as log-transformed number of ITS copies per μl of reaction mix) in the soil of the experimental vineyard (top right panel; open circles size is proportional to the number of data pairs; solid line corresponds to the fitted Matern model). Map of the experimental vineyard with interpolated values of *P. viticola* DNA (expressed as the number of ITS copies per μl of reaction mix) obtained by ordinary kriging (left bottom panel). Map of the kriging standard deviation (right bottom panel).

The pattern of the plot-scale experimental semivariogram was not typical, with semivariance alternatively increasing and decreasing along the gradient of separation distance (Figure S8 in Supplementary Materials). Semivariance peaks corresponded to distance classes with a higher proportion of heterogeneous pairs of points, *i.e.* pairs consisting of one sampling point in a row and one sampling point in an inter-row. On the other hand, lowest semivariance values corresponded to distance classes with a higher proportion of homogeneous pairs of points, *i.e.* pairs with two sampling points in a row or pairs with two sampling points in an inter-row (Figure S8 in Supplementary Materials). We consistently found that semivariance was higher for heterogeneous pairs of points than for homogeneous ones, on average (anova P <0.0001, Figure S8 and Table S6 in Supplementary Materials).

### *Plasmopara viticola* primary inoculum along depth and distance-to-row gradients

Our results clearly indicated that there was a decreasing gradient of primary inoculum from the row to the middle of the inter-row (Figure 3; β = −0.0263 ± 0.0023; df = 133; t value = −11.54; P < 0.001; R²m = 0.48; R²c = 0.50). Downy mildew primary inoculum was on average five to six times more concentrated in the ridge of soil below the vine stocks (9.32 ± 6.32 ITS copies per µl) than in the inter-row space (1.72 ± 1.66 ITS copies per µl). These results were in line with the results from the geostatistical analysis of nested sampling plot data (see above). We also found that the primary inoculum concentration decreased with soil depth (Figure 3; Kruskal-Wallis test’s statistic = 12.1; df = 3; P < 0.01). In particular, it was significantly lower between 21 and 30 cm and between 31 and 40 cm than in the first 10 cm from the soil surface (see results of Dunn’s post-hoc test in Table S7 in Supplementary Materials).

**Figure 3.**
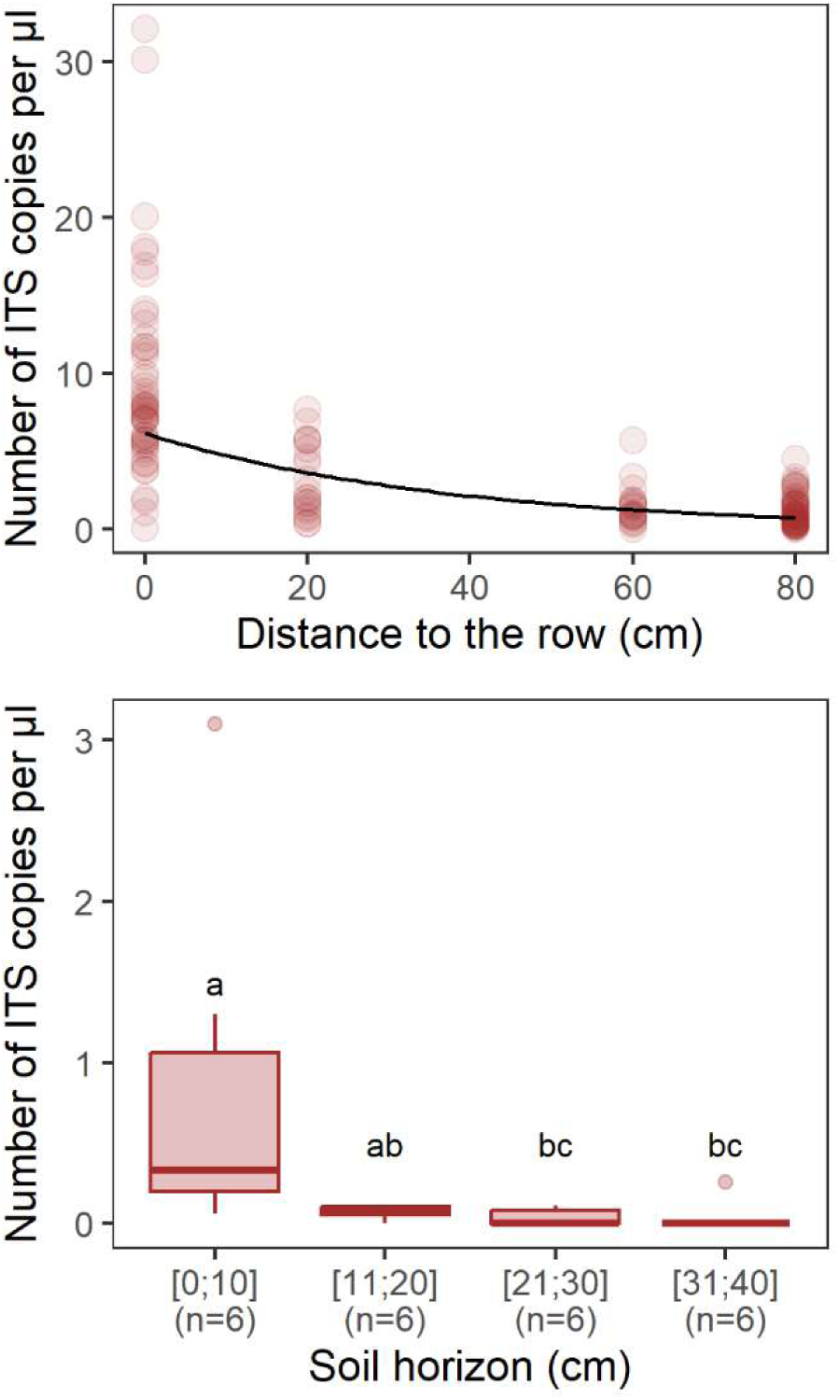
Abundance of downy mildew primary inoculum (as number of ITS copies per µl estimated by ddPCR) along gradients of distance to the row (upper panel) and soil depth (lower panel). Solid line in the upper panel corresponds to model predictions. Different letters between depths in the lower panel indicate a significant difference.

Although we were not seeking to address this point, we also found that primary inoculum concentration in the soil varied between rows, with significantly less *P. viticola* DNA on average in the westernmost row compared to others in the vineyard (Figure S9 Supplementary Materials; F = 6.522; P < 0.001; see Table S8 in Supplementary Materials for results of Tukey’s multiple comparison test). This unexpected result was consistent with the distribution of primary inoculum as predicted by kriging of observed data (Figure 2).

### A bioassay to assess soil infectious potential

The cumulated number of leaf discs infected by *Plasmopara viticola* from 1g of soil sample over the duration of the assay varied from 0 to 12 discs (out of 40). The average proportion of disc area covered with *P. viticola* sporulation varied from 0 to 10.3%. The cumulated number of infected discs significantly increased with the number of ITS copies per µl (Figure 4; ꞵ = 0.029 ± 0.008; P < 0.0001). In addition, the average infected area significantly increased with the number of ITS copies per µl (Figure 4; ꞵ = 0.058 ± 0.010; P < 0.0001). The coefficients of determination were low in both cases, although it was slightly higher for the average infected area than for the cumulated number of infected discs (R² = 0.33 and 0.19, respectively).

**Figure 4.**
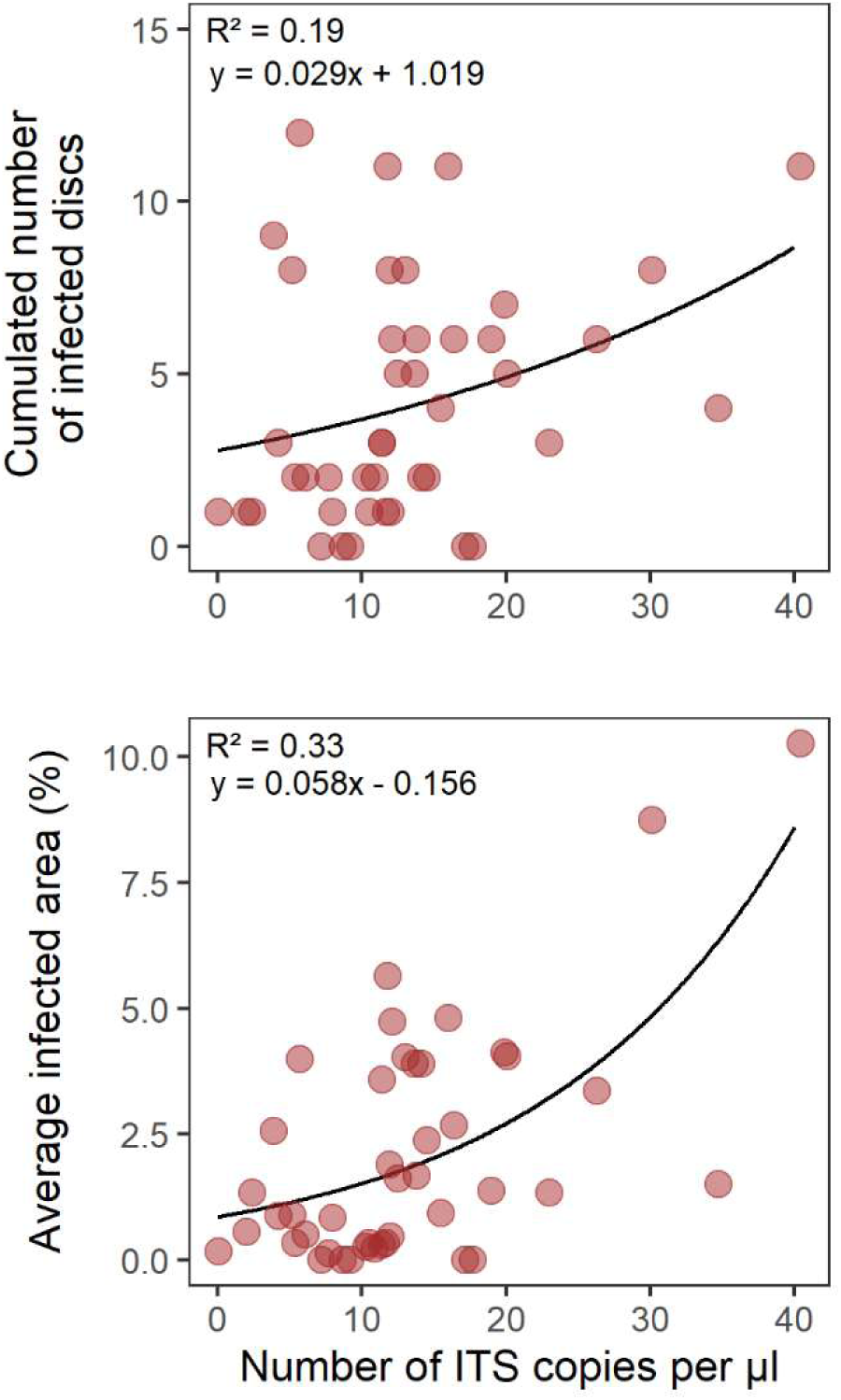
Comparison of bioassay and digital droplet PCR. Relationships between the abundance of DM primary inoculum in the soil (measured as number of ITS copies per µl by ddPCR) and the infectious potential of the soil measured by a bioassay, as either the cumulated number of infected discs (upper panel) or the average trap disc infected area (lower panel). Solid lines represent fitted model predictions.

## Discussion

We presented a method for the quantification of *P. viticola* oospores in vineyard soil samples based on a ddPCR approach. We used this method to describe the spatial distribution of downy mildew primary inoculum (oospores) at different spatial scales in one vineyard. We also evaluated whether the ddPCR-based assessment of inoculum concentration was related to the soil infectious potential as assessed using a bioassay. The method used for inoculum assessment in soil is based on the ddPCR technique, which provides an absolute quantification of DNA and is particularly well suited for rare targets in complex matrices. We combined these advantages to the use of an ITS sequence occurring in numerous copies which increases the probability of detection of *P. viticola* DNA in small quantities in the sample. The partition of the samples into thousands of droplets improves the yield of amplification and the sensitivity of the analysis as compared to qPCR, including in the case of oomycete quantification in soil samples (Gossen *et al*., 2019; Rački *et al*., 2014; Wen *et al*., 2020). Using ddPCR and the average estimate for the number of ITS1/5.8S copy in the genome of *P. viticola* (215 copies), we found that the number of oospores per gram of soil ranged from 0 to 1858 in the experimental vineyard and averaged 303 oospores per gram of soil. This result falls within the range of values (*i.e.* from a few dozen to 4000 oospores per gram of soil approximately) previously estimated by Yang *et al*. (2023) in Chinese vineyards using qPCR.

The analysis of soil samples spiked with known concentrations of oospores revealed that our method can detect *P. viticola* oospore DNA at very low concentration (*i.e.* 1 oospore in 2 g of soil) and is an adequate and repeatable tool for the relative quantification of *P. viticola* oospore DNA in vineyard soil. However, the method showed some limitations for absolute quantification purpose, in particular because of variable (and sometimes low) efficiency and underestimation bias. Such limitations reflect the accumulation of errors regarding the representativeness of sampling after soil homogenization, technical errors during sample preparation, DNA extraction (*e.g.* irregular pipetting) or ddPCR assay. Further methodological improvements such as automation could help reduce them.

A potential drawback of the use of a multicopy DNA sequence as a primer lies in the potential variability of the number of copy of the sequence among genotypes. In our study, we found that the number of ITS1/5.8S copies in *P. viticola* diploid genome varied from 175 to 281 copies depending on the strain (n=7), which caused the primary inoculum concentration estimates to vary from 232 to 372 oospores per gram of soil on average. The range of ITS copy number found in our study is included in the range estimated for the oomycete *Aphanomyces euteiches*, but is more restricted (meaning it is probably underestimated), probably because it is estimated from a smaller number of strains (Gangneux *et al*., 2014; Gibert *et al*., 2021). Further analyses should be undertaken to better assess the occurrence of the ITS1/5.8S on a *P. viticola* population level.

Finally, *P. viticola* may persist under various forms in vineyard soils, including oospores and non-viable structures such as dead mycelial cells from fallen leaves, as well as sporangia or zoospores that settle on the ground without encountering a host. Consequently, DNA-based methods used to quantify downy mildew inoculum reservoir in vineyard soils must be interpreted with caution as extracellular DNA could be amplified, leading to an overestimation of the viable inoculum. In our study, soil sampling was deliberately conducted well after the end of the growing season (mid-March) to maximize extracellular DNA degradation and to limit this bias. In microcosm experiment, bacterial and plant extracellular DNA were found to persist from a few hours to 70 days and from 3 months to 2 years in the soil, respectively (Nielsen *et al*., 2007). The rate of degradation varied depending on abiotic conditions, soil management practices and whether the DNA was protected from enzymatic degradation within plant tissue (Nielsen *et al*., 2007; Sirois and Buckley, 2019). No data have been published on oomycete extracellular DNA. To further assess and reduce the risk of overestimating the viable inoculum, methods that prevent extracellular DNA amplification should be considered, such as treating the samples with propidium monoazide (PMA) before DNA extraction and/or reverse-transcriptase qPCR (see for example Wang *et al*., 2023). Such approaches have already proven to improve DNA-based quantitative assessment of viable plant pathogens or plant-product contaminants (Al-Daoud *et al*., 2017; Crespo-Sempere *et al*., 2013).

The relatively high nugget value found in the semivariogram (0.156) indicated that very close points could be quite dissimilar in terms of primary inoculum, suggesting the existence of a source of variation operating at a scale finer than that of the regular grid (*i.e.* sub-grid variability). Alternatively, it could reflect measurement error related to the methodological limitations in the ddPCR approach discussed above (*i.e.* ITS copy number variability, amplification of extracellular DNA, variable efficiency). In our study, *P. viticola* was encountered at all sampled locations in the field contrary to soilborne pathogens whose presence is generally restricted to some areas (Botero-Ramirez *et al*., 2021; Moussart *et al*., 2009). This difference may be explained by the greater dispersal ability of the airborne form of *P. viticola* (as compared to strictly soilborne pathogens) resulting in a more even distribution of the disease in the field. Still, we showed that the distribution of downy mildew oospores was not random or regular on a field scale, but that it was characterized by 15 m-diameter patches of concentrically increasing oospore concentration. This pattern could be related to the distribution of the disease in the vineyard at the end of the growing season, which we unfortunately did not assess the year before our soil survey (but see Box S3 in Supplementary materials for comparisons of the spatial distribution of primary inoculum against soil electrical resistivity and grapevine NDVI distributions, as indirect proxies for disease incidence). Alternatively, environmental factors could affect grapevine leaf litter (containing the oospores) distribution or the survival or maturation of the inoculum during autumn and winter (*e.g.* copper or other fungicidal substances, soil moisture, antagonistic microbial species presence). Future studies should investigate the relationship between the spatial distribution of the symptoms at the end of the growing season and the distribution of the inoculum in the soil the following year.

We found that, on average, oospores accumulates preferentially in the ridge of soil below the vine stocks (or “row location”) at a rate of 4 to 5 times the inter-row space, and in the first 10 cm of soil. Although oospore concentration varied on a wider spatial scale within the field (15m patches), this row *vs* inter-row contrast was observed at all locations in the field. Moreover, oospore concentration was more similar at two row locations, whatever the distance between the two points, than when comparing row *vs* inter-row locations. This suggests that the distribution of oospores is primarily determined by the distribution of grapevine leaf litter that carry them, which falls and accumulates at the foot of the grapevines. However, the distribution of grapevine leaf litter may change during autumn and winter under the influence of the wind and of cultural practices. Instead, we hypothesize that soil ridging at the foot of the vines has reinforced this distribution pattern by preventing the dispersion of the fallen leaves by the wind during winter (see Figure S10 in Supplementary Materials for an illustration)

Our study also revealed that soil oospore concentration can vary between rows within a field and be subjected to border effects. Indeed, we found that the primary inoculum was 7 to 8 times less concentrated in the westernmost row compared to the rest. This difference might arise from various mechanisms mediated by the presence of a tree hedgerow alongside the experimental field, possibly limiting disease incidence of leaf litter abundance locally. These mechanisms include i) wind erosion and microclimate modification in the vicinity of the hedgerow (Campi *et al*., 2009; Jacobs *et al*., 2022), ii) modulation of grapevine vigor and susceptibility to downy mildew potentially due to competition for resources (Favor and Udawatta, 2021; but see Bourgade *et al*., 2020), and iii) interception of airborne spores (Delatouche *et al*., 2024). In our case, the absence of a border effect on the east side of the field (also bordered by a hedgerow) suggests that mechanisms at play operate asymmetrically, which restricts the choice to mechanisms associated to wind erosion and microclimate modification. In any case, these results raise questions regarding the effects of agroecological infrastructures on downy mildew epidemiology, as well as their possible contribution to preventive disease management strategies.

Positive relationships between inoculum abundance and disease severity were found for soilborne root pathogens (Chatterton *et al*., 2023; Hwang *et al*., 2011; Moussart *et al*., 2009), but the inoculum density-disease severity relationship has rarely been studied for soil-to-aerial plant parts infection pathway, such as in the case of downy mildew primary contaminations. Only one recent study demonstrated that the risk of primary infections of grapevine leaves by *P. viticola* increased with the concentration of oospores in the leaf litter deposited at the foot of the vine stocks (Fedele *et al*., 2025), but the experimental conditions were not particularly representative of the amount of primary inoculum reservoir found “naturally” in vineyards (important amount of infected leaf litter artificially accumulated at the foot of the grapevines). Our results supported the idea that the concentration of *P. viticola* oospores in the soil is related to soil infectious potential (as estimated in controlled conditions using a leaf disc bioassay). In particular, we found that the proportion of leaf discs infected during the bioassay significantly increased with primary inoculum abundance in the soil (as estimated by ddPCR). However, the relationship was not so strong (R² = 0.33) which was possibly due to the limitations in DNA-based estimation of primary inoculum reservoir discussed above. Moreover, as oospore germination in a population extends over several months (Maddalena *et al*., 2021), our bioassay (carried out on a single date) likely “captured” only part of the oospore population present in the samples. Besides technical limitations, overlooked factors driving oospore germination could contribute to lower the strength of the relationship that we found. First, oospores can survive and germinate at least for five years, meaning that oospores of different ages coexist in a vineyard soil at time t (Caffi *et al*., 2011). The effect of aging on oospore germination remains undescribed, but it is likely that the outcome of the germination bioassay will depend on the age pyramid of the oospores present in the sample. In addition, autumn and winter conditions can affect oospore germination rate and timing (*e.g.* Jermini *et al*., 2003; Rouzet and Jacquin, 2003), so that spatial heterogeneity of conditions (including within a field) can lead to variability in the outcome of the germination bioassay between samples with similar *P. viticola* oospores or DNA contents. Altogether, this suggests that, even if the infectious potential is undoubtedly related to the amount of primary inoculum (which can be assessed by means of a DNA-based approach) to some extent, the indicators derived from our germination bioassay should be complemented with contextual data for epidemiological interpretation.

As agriculture moves away from chemical biocides, the design of sustainable, adaptative and cost-effective crop disease management strategies will rely more and more on the prevention of epidemic risks and a predictive evaluation of pathogen population dynamics. In this context, the quantitative and spatially-explicit survey of inoculum reservoirs is crucial. The replication of primary inoculum survey in different contexts and years, with varying pedoclimatic, epidemiological or cultural conditions, will provide a more generic understanding of the abundance and spatial distribution of *P. viticola* primary inoculum in vineyards. Altogether, a better characterization of the variability of ITS1/5.8S copy number among *P. viticola* genotypes, the consideration of extracellular *P. viticola* DNA in vineyard soils and methodological improvements to limit measurement error will enable more accurate estimates of the quantity of primary inoclum in the soil and a better understanding of the relationship between the quantity of inoculum and epidemics. Ultimately, DNA-based survey of primary inoculum could contribute to the design of monitoring and decision-making tools (*e.g.* epidemic risk prediction models) to accompany the implementation of sanitation measures preventing primary inoculum accumulation in the vineyard from one season to the next.

## Acknowledgements

The authors thank the Experimental Viticultural Unit of Bordeaux 1442, INRAE, F-33883 Villenave d’Ornon, for the maintenance of the experimental vineyard that was used in this study, and the GenoSol platform (DOI 10.15454/L7QN45) for sharing its expertise and helping with methodological development for DNA extraction from soil samples. Part of the experiments (ddPCR analysis) were performed at the Genome Transcriptome Platform of Bordeaux (PGTB; doi:10.15454/1.5572396583599417E12) with the help of Adline Delcamp and Emilie Chancerel. Finally, the authors are grateful to the sponsors of the Chaire Alexis Millardet for their support to research activities, namely Château Ausone, Château Cheval Blanc, Petrus and Château Montrose.

