## Supplementary materials for "Direct quantitative assessment using digital droplet PCR and field-scale spatial distribution of *Plasmopara viticola* oospores in vineyard soil"

Poeydebat *et al.*

**Box S1. Method for oospore production**

Four *P. viticola* strains of compatible mating types (two P1 and two P2 strains) were asexually multiplied, separately, on Cabernet sauvignon leaves in Petri dishes at 20°C and the sporangia of the four strains were used to prepare one suspension containing the four strains at equal concentrations.

We collected leaves (the eighth from the apex) from Cabernet sauvignon grapevines grown in experimental greenhouses and prepared 16mm diameter leaf discs that we deposited abaxial face up in Petri dishes filled with 20% agar. Leaf discs were co-infected with the four *P. viticola* strains with compatible mating types by depositing droplets of the suspension (three 15µl-droplets per disc), and then left at 20°C for 7 days until *P. viticola* sporulated. At this time, we placed the discs at 10°C for 3 weeks, which triggered sexual reproduction between strains of compatible mating types. After 3 weeks, the presence of oospores in the leaf discs was checked using a microscope, and the discs were placed at 5°C for at least 2 months to allow for oospore maturation.

Then leaf discs were gently grinded with a porcelain pestle and mortar in a few mL of distilled water and the obtained suspension was filtered at successively 100, 60 and 40µm to separate the oospores from leaf disc fragments. The filtrate was finally passed through a 20µm filter that retained most oospores, and the oospores were re-suspended in a small volume of distilled water. Oospore concentration of the final suspension was estimated using a hematocytometer (5 count replicates). This suspension was used to inoculate soil samples at known concentrations.

**Table S1.** Serial dilution of *Plasmopara viticola* oospores in non-infected 2g-soil samples, used for determining the linearity and efficiency of ddPCR. The expected number of ITS copies per µl of reaction mix was computed on the basis of the number of oospores added in the initial sample and the average number of ITS (1/5.8S) copies in the diploid genome of *P. viticola* estimated in this study (*i.e.* 215 copies).

| Number of oospores in 2g of soil | Number of ITS copies per µl of reaction mix | | Efficiency (%) | Number of replicates | Detection frequency (rate of positive replicates) |
|---|---|---|---|---|---|
| | Expected | ddPCR result (mean ± sd) | | | |
| 1 | 0.011 | 0.010 ± 0.022 | 92 | 5 | 1/5 |
| 10 | 0.109 | 0.012 ± 0.027 | 11 | 5 | 1/5 |
| 50 | 0.543 | 0.316 ± 0.089 | 58 | 5 | 5/5 |
| 100 | 1.087 | 0.360 ± 0.168 | 33 | 5 | 5/5 |
| 500 | 5.435 | 2.340 ± 0.991 | 43 | 5 | 5/5 |
| 1000 | 10.869 | 8.800 ± 1.570 | 81 | 5 | 5/5 |

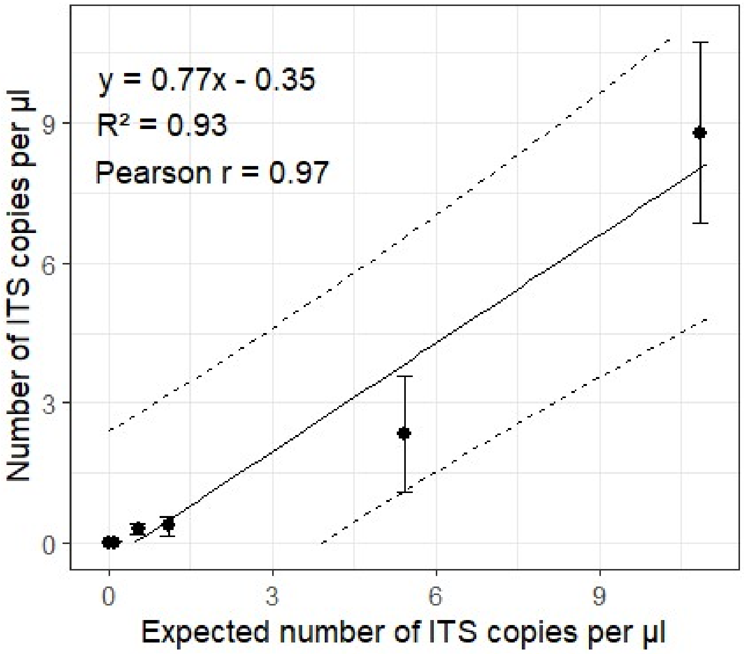


**Figure S1.** Linear regression analysis of the ddPCR assay based on a serial dilution of *Plasmopara viticola* oospores in non-infected 2g-soil samples. The vertical axis corresponds to the number of ITS copies per µl of reaction mix estimated in the ddPCR assay. The horizontal axis corresponds to the expected number of ITS copies per µl of reaction mix, based on the number of oospores added in the initial sample and the average number of ITS (1/5.8S) copies in the genome of *P. viticola* estimated in this study (*i.e.* 215 copies). The error bars represent the 95% confidence interval around the mean based on five replicates. The solid and dotted lines are the linear regression and 95% confidence interval, respectively.

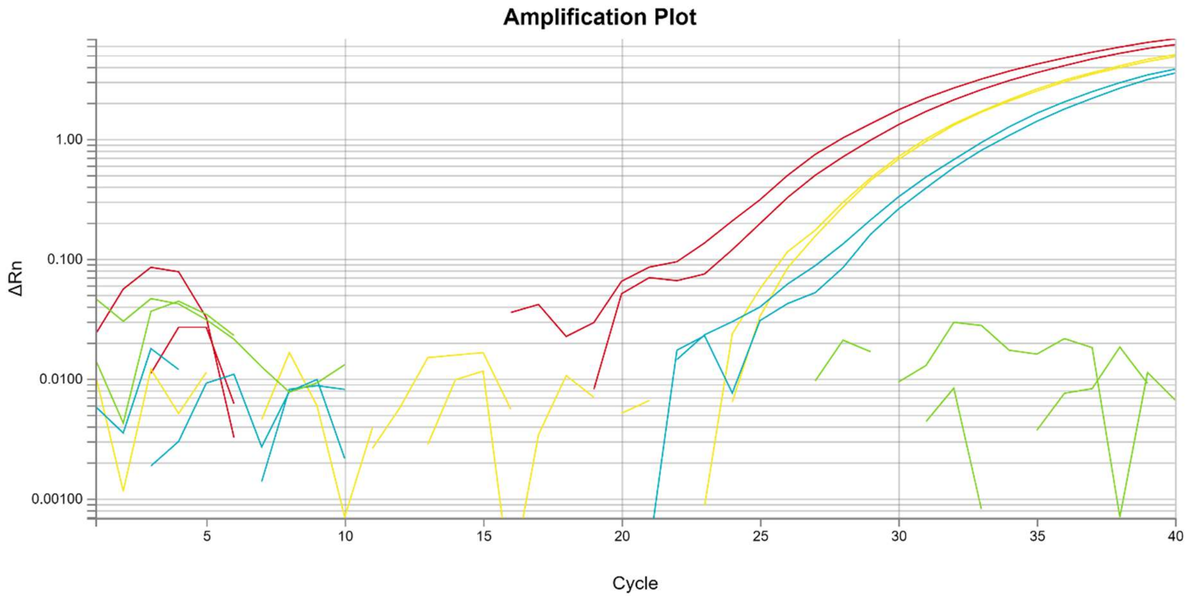


**Figure S2.** Amplification plot of the qPCR analyses testing for β-tubulin primers specificity for *Plasmopara viticola*. Red: Pure *P. viticola* strain diluted to the tenth. Yellow: Pure P. viticola strain diluted to the hundredth. Blue: *P. viticola* + grapevine leaf. Green: grapevine leaf.

**Table S2.** ITS and β-tubulin copy number obtained by ddPCR for several *Plasmopara viticola* (Pv) strains.

| Pv strain (or $H_2O$) | DNA (ng) | Target | Concentration | Copies/ng | Number of copy in the diploid genome |
|---|---|---|---|---|---|
| **$H_2O$** | 0 | B_Tub | No Call | N/A | N/A |
| **3174_11** | 5 | B_Tub | 732 | 146.4 | 1 |
| **221** | 5 | B_Tub | 2018 | 403.6 | 1 |
| **412_11** | 5 | B_Tub | 1516 | 303.2 | 1 |
| **2664** | 5 | B_Tub | 1457 | 291.4 | 1 |
| **2543** | 5 | B_Tub | 1539 | 307.8 | 1 |
| **8676** | 5 | B_Tub | 1837 | 367.4 | 1 |
| **8239** | 5 | B_Tub | 755 | 151 | 1 |
| **$H_2O$** | 0 | ITS | No Call | N/A | N/A |
| **3174_11** | 0.005 | ITS | 206 | 41200 | 281 |
| **221** | 0.005 | ITS | 465 | 93000 | 230 |
| **412_11** | 0.005 | ITS | 364 | 72800 | 240 |
| **2664** | 0.005 | ITS | 282 | 56400 | 194 |
| **2543** | 0.005 | ITS | 270 | 54000 | 175 |
| **8676** | 0.005 | ITS | 345 | 69000 | 188 |
| **8239** | 0.005 | ITS | 150 | 30000 | 199 |

**Table S3.** Settings of the plot-scale experimental omnidirectional semivariogram.

| Distance lag (m) | Maximum distance (m) | Total number of point pairs | Number of distance classes | Number of point pairs per distance class | |
|---|---|---|---|---|---|
| | | | | min | max |
| 0.2 | 4 | 1863 | 20 | 30 | 206 |

| Lag class | Bounds (m) | | Number of pairs |
|---|---|---|---|
| | Lower | Upper | |
| 1 | 0 | 0.2 | 62 |
| 2 | 0.2 | 0.4 | 74 |
| 3 | 0.4 | 0.6 | 124 |
| 4 | 0.6 | 0.8 | 206 |
| 5 | 0.8 | 1.0 | 161 |
| 6 | 1.0 | 1.2 | 48 |
| 7 | 1.2 | 1.4 | 38 |
| 8 | 1.4 | 1.6 | 50 |
| 9 | 1.6 | 1.8 | 76 |
| 10 | 1.8 | 2.0 | 62 |
| 11 | 2.0 | 2.2 | 117 |
| 12 | 2.2 | 2.4 | 135 |
| 13 | 2.4 | 2.6 | 121 |
| 14 | 2.6 | 2.8 | 131 |
| 15 | 2.8 | 3.0 | 99 |
| 16 | 3.0 | 3.2 | 125 |
| 17 | 3.2 | 3.4 | 89 |
| 18 | 3.4 | 3.6 | 40 |
| 19 | 3.6 | 3.8 | 50 |
| 20 | 3.8 | 4 | 30 |

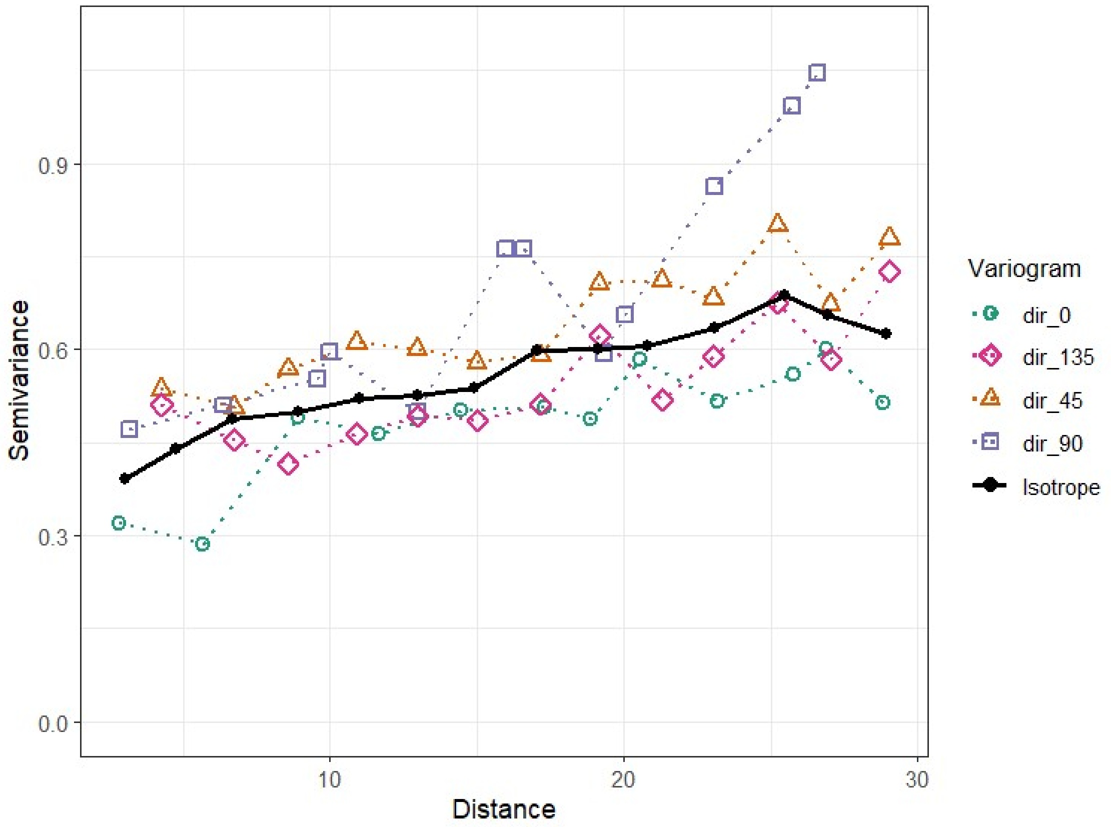


**Figure S3.** Omnidirectional (isotrope) and directional (0°, 45°, 90° and 135°) experimental semivariograms of *P. viticola* DNA in soil (expressed as log-transformed number of ITS copies per µl of reaction).

**Table S4**. Structural parameters of the experimental directional semivariograms of *P. viticola* DNA in soil (expressed as log-transformed number of ITS copies per µl of reaction).

| Direction | Range | Sill | Nugget |
|---|---|---|---|
| 0° | 19.849 | 0.288 | 0.250 |
| 45° | 4.285 | -0.064 | 0.688 |
| 90° | 200.002 | 1.612 | 0.459 |
| 135° | 6.461 | -0.293 | 0.812 |

**Table S5.** Output of the model selection procedure based on the autofitVariogram() function of the R package 'autofit'. Theoretical models were fitted to the experimental omnidirectional semivariogram of *P. viticola* DNA in soil (expressed as log-transformed number of ITS copies per µl of reaction mix).

| Model | Sum of squared errors (sserr) |
|---|---|
| Spherical | 0.0317 |
| Matérn (kappa=0.05) | 21.8691 |
| Matérn (kappa=0.2) | 0.0297 |
| Matérn (kappa=2) | 0.0406 |
| Matérn (kappa=10) | 0.0510 |
| Exponential | 0.0306 |

**Figure S4.** Raw and detrended (linear or quadratic trend) omnidirectional experimental semivariograms of *P. viticola* DNA in soil (expressed as log-transformed number of ITS copies per µl of reaction mix).

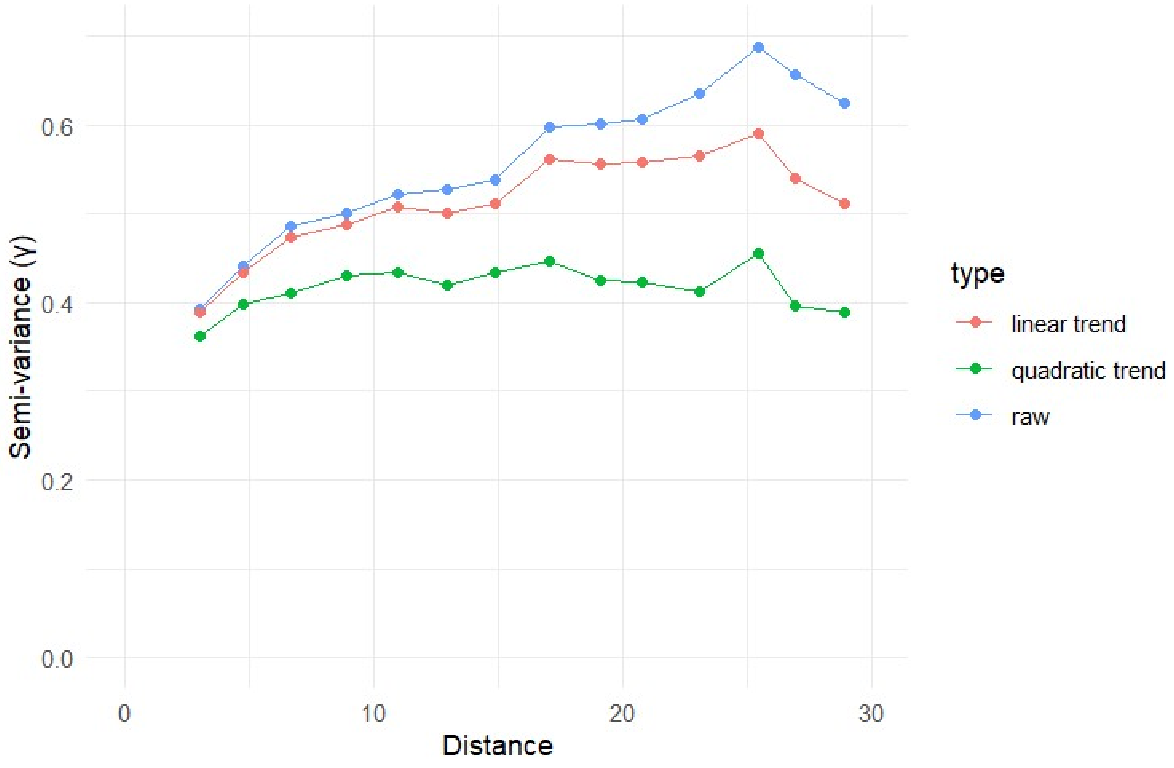

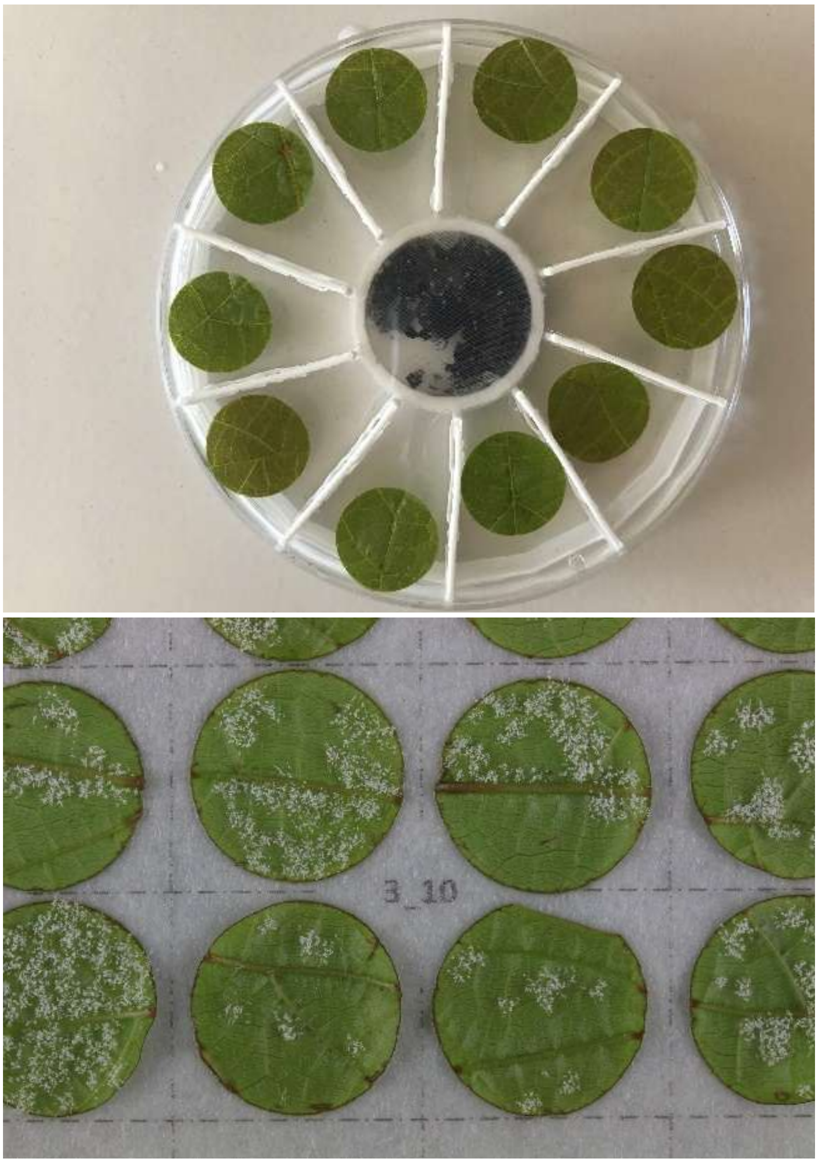


**Figure S5.** Photographies of the floating leaf disc bioassay. On top, the photo shows the circular box with the leaf disc separation structure, the ten grapevine leaf discs floating on water and the soil sample in the middle of the box held at the bottom with the blotting cloth. At the bottom, the photo shows the incubated leaf discs covered with *Plamsopara viticola* sporulation after infection by zoospores released by oospores.

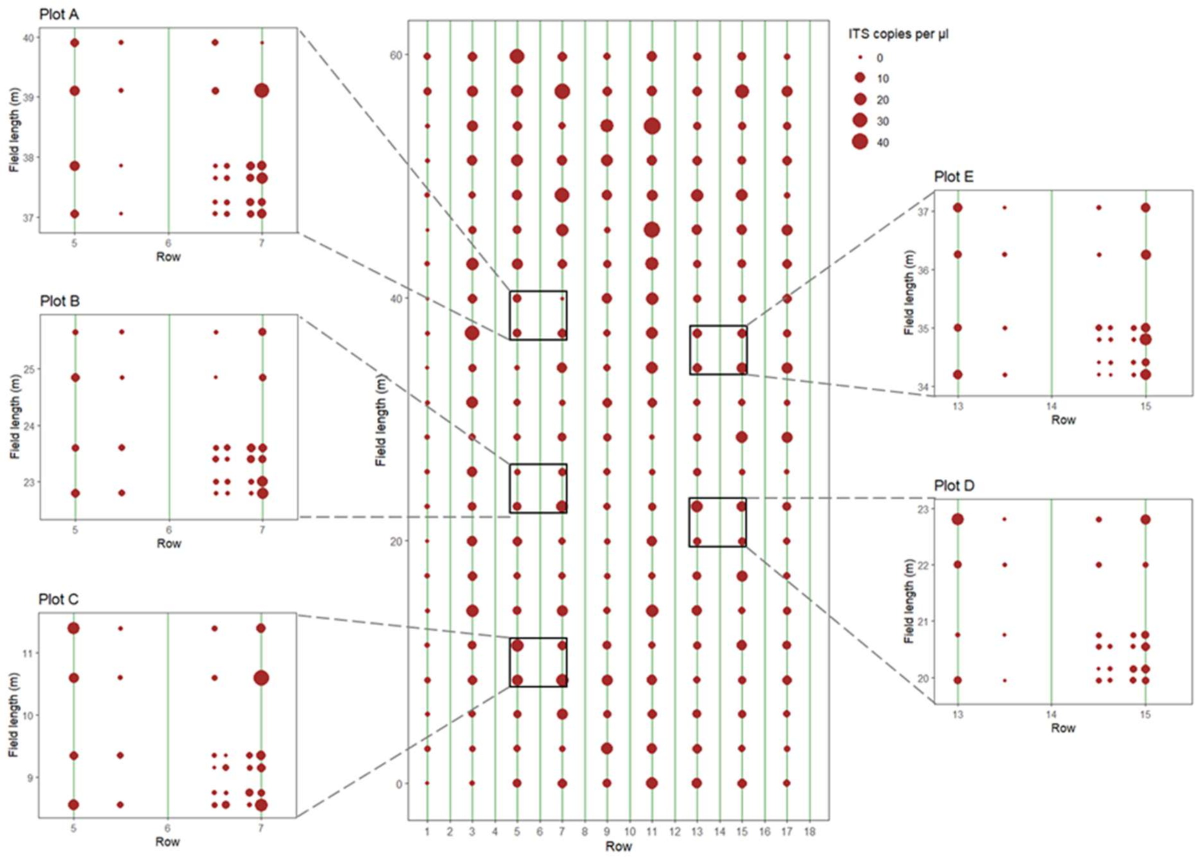


**Figure S6.** Top view of the experimental vineyard and spatial distribution of *Plasmopara viticola* primary inoculum (as number of ITS copies per µl of reaction mix) found in the soil, in samples of both the regular grid and the nested sampling plots A to E.

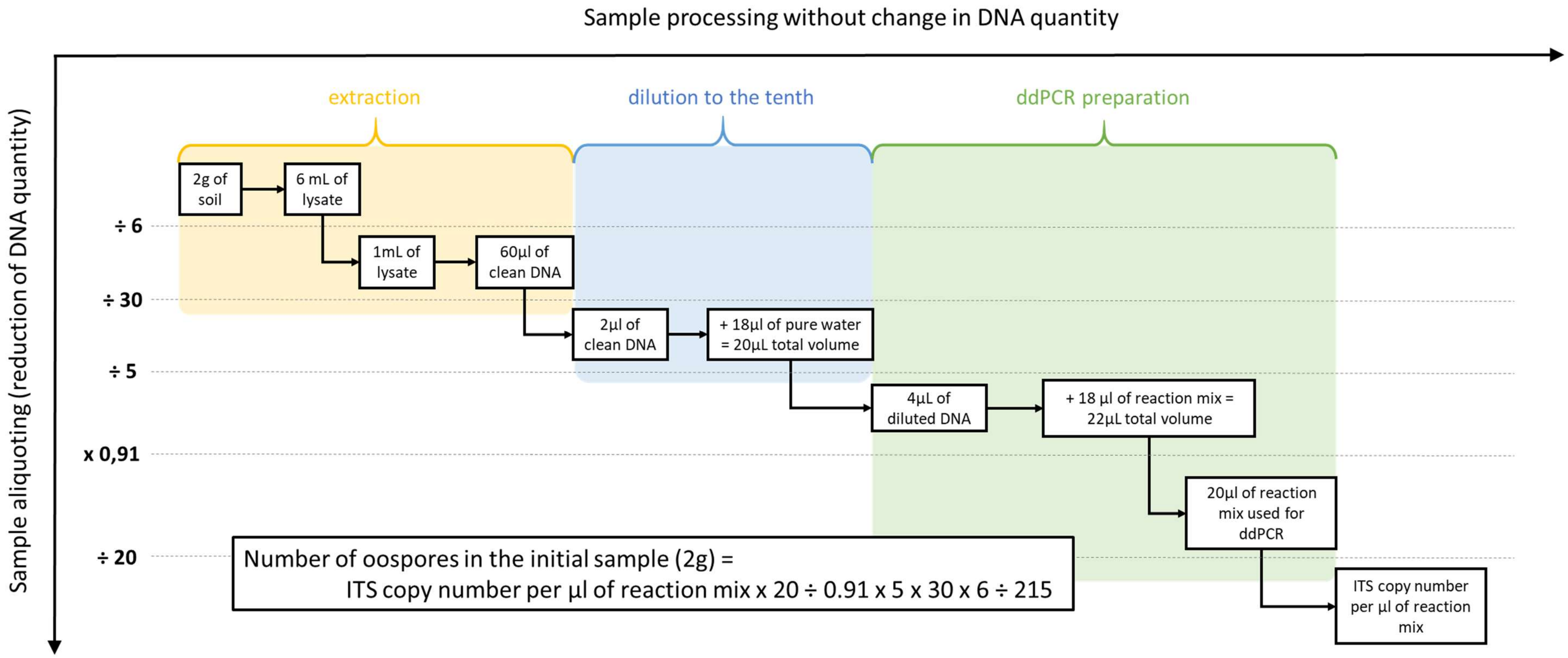


**Figure S7.** Schematic representation of the DNA extraction and ddPCR processing and reverse chain of computation to convert the number of ITS1/5.8S copies per µl of reaction mix into the number of oospores in the initial sample (2g of soil). We assumed that the extraction process was 100% efficient (100% of DNA recovery). The number 215 corresponds to the number of ITS1/5.8S (primers sequence) copies in the genome of *Plasmopara viticola*. It was obtained from the comparison of results of ddPCR based on ITS *vs* β-tubulin (single copy) primers, averaged over 7 *Plasmopara viticola* (Pv) strains (see Table S2).

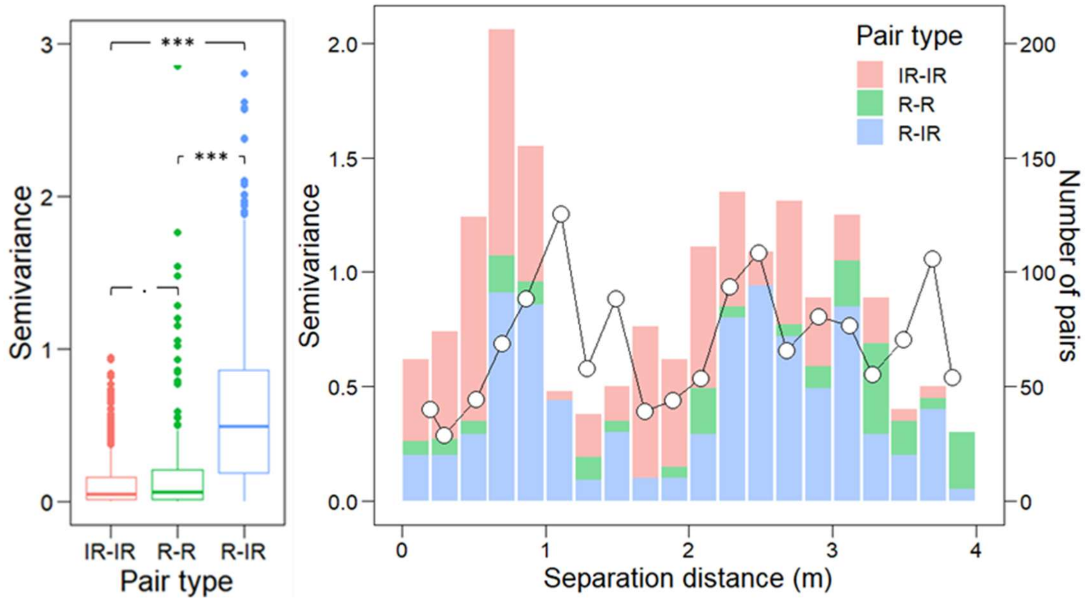


**Figure S8.** Mean semivariance of *P. viticola* DNA (expressed as log-transformed number of ITS copies per µl of reaction mix) in soil per type of sampling point pairs (left panel). Plot-scale experimental semivariogram of *P. viticola* DNA (expressed as log-transformed number of ITS copies per µl of reaction mix) in soil and number of point pairs per distance class and per type of pair (right panel). IR: inter-row; R: row.

**Table S6.** Tukey's multiple comparison test for *P. viticola* DNA (expressed as log-transformed number of ITS copies per µl of reaction mix) semivariance per type of point pair (nested sampling plot data). R: row; IR: inter-row. R-R and IR-IR are considered homogeneous pairs of points, while R-IR are heterogeneous pairs of points

| Single comparison (type of pair) | Test scores difference | 95% confidence interval | | Adjusted p-value |
|---|---|---|---|---|
| | | lower bound | upper bound | |
| R-R vs IR-IR | 0.065 | -0.002 | 0.133 | 0.058 |
| R-IR vs IR-IR | 0.469 | 0.425 | 0.513 | **0.000** |
| R-IR vs R-R | 0.403 | 0.337 | 0.469 | **0.000** |

**Table S7**. Dunn's multiple comparison test for the downy mildew primary inoculum at different depths in the soil. P-values in bold indicate a significant difference (at a threshold of 0.05) in primary inoculum concentration between two depths.

| Single comparison (depths) | Dunn test's statistic | Adjusted p-value |
|---|---|---|
| [0;10] vs [11;20] | -1.89 | 0.354 |
| [0;10] vs [21;30] | -2.86 | **0.025** |
| [0;10] vs [31;40] | -3.14 | **0.010** |
| [11;20] vs [21;30] | -0.98 | 1 |
| [11;20] vs [31;40] | -1.25 | 1 |
| [21;30] vs [31;40] | -0.28 | 1 |

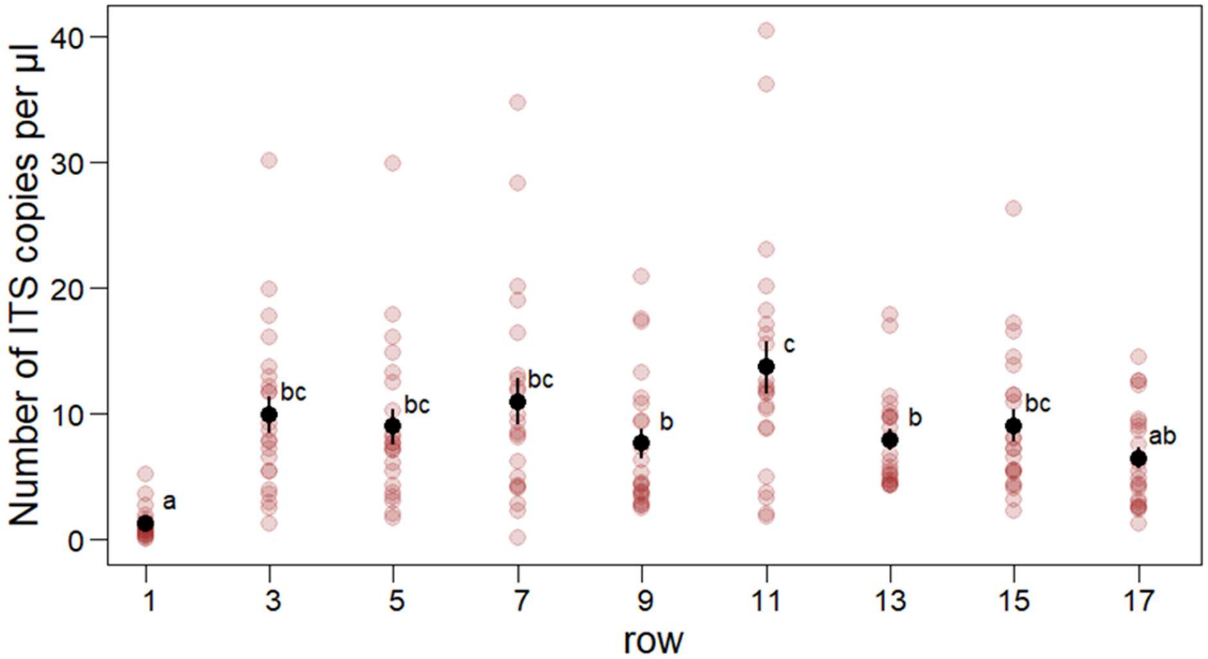


**Figure S9.** Downy mildew primary inoculum (as number of ITS copies per µl of reaction mix) in different vine stock rows of the experimental vineyard. Rows are numbered from 1 to 17 from west to east. Red dots correspond to observed values. Black dots and error bars correspond to mean and standard errors, respectively. Different letters indicate significant differences between rows.

**Table S8.** Tukey's multiple comparison test for the downy mildew primary inoculum in different vine stock rows within the surveyed experimental vineyard. Rows are numbered from 1 to 17 from west to east. P-values in bold indicate a significant difference (at a threshold of 0.05) in primary inoculum concentration between two rows.

| Single comparison (rows) | Test scores difference | 95% confidence interval | | Adjusted p-value |
|---|---|---|---|---|
| | | lower bound | upper bound | |
| 3-1 | 8.6968 | 2.7481 | 14.6456 | **0.0003** |
| 5-1 | 7.7314 | 1.7826 | 13.6801 | **0.0021** |
| 7-1 | 9.7341 | 3.7853 | 15.6829 | **0.0000** |
| 9-1 | 6.3677 | 0.4190 | 12.3165 | **0.0259** |
| 11-1 | 12.4632 | 6.5144 | 18.4119 | **0.0000** |
| 13-1 | 6.6723 | 0.7235 | 12.6210 | **0.0155** |
| 15-1 | 7.8223 | 1.8735 | 13.7710 | **0.0018** |
| 17-1 | 5.2255 | -0.7233 | 11.1742 | 0.1357 |
| 5-3 | -0.9655 | -6.9142 | 4.9833 | 0.9999 |
| 7-3 | 1.0373 | -4.9115 | 6.9860 | 0.9998 |
| 9-3 | -2.3291 | -8.2779 | 3.6197 | 0.9492 |
| 11-3 | 3.7664 | -2.1824 | 9.7151 | 0.5545 |
| 13-3 | -2.0245 | -7.9733 | 3.9242 | 0.9780 |
| 15-3 | -0.8745 | -6.8233 | 5.0742 | 0.9999 |
| 17-3 | -3.4714 | -9.4201 | 2.4774 | 0.6614 |
| 7-5 | 2.0027 | -3.9460 | 7.9515 | 0.9795 |
| 9-5 | -1.3636 | -7.3124 | 4.5851 | 0.9985 |
| 11-5 | 4.7318 | -1.2169 | 10.6806 | 0.2406 |
| 13-5 | -1.0591 | -7.0079 | 4.8897 | 0.9998 |
| 15-5 | 0.0909 | -5.8579 | 6.0397 | 1.0000 |
| 17-5 | -2.5059 | -8.4547 | 3.4429 | 0.9237 |
| 9-7 | -3.3664 | -9.3151 | 2.5824 | 0.6980 |
| 11-7 | 2.7291 | -3.2197 | 8.6779 | 0.8809 |
| 13-7 | -3.0618 | -9.0106 | 2.8869 | 0.7955 |
| 15-7 | -1.9118 | -7.8606 | 4.0369 | 0.9847 |
| 17-7 | -4.5086 | -10.4574 | 1.4401 | 0.3020 |
| 11-9 | 6.0955 | 0.1467 | 12.0442 | **0.0400** |
| 13-9 | 0.3045 | -5.6442 | 6.2533 | 1.0000 |
| 15-9 | 1.4545 | -4.4942 | 7.4033 | 0.9976 |
| 17-9 | -1.1423 | -7.0910 | 4.8065 | 0.9996 |
| 13-11 | -5.7909 | -11.7397 | 0.1579 | 0.0631 |
| 15-11 | -4.6409 | -10.5897 | 1.3079 | 0.2646 |
| 17-11 | -7.2377 | -13.1865 | -1.2890 | **0.0056** |
| 15-13 | 1.1500 | -4.7988 | 7.0988 | 0.9996 |
| 17-13 | -1.4468 | -7.3956 | 4.5019 | 0.9977 |
| 17-15 | -2.5968 | -8.5456 | 3.3519 | 0.9077 |

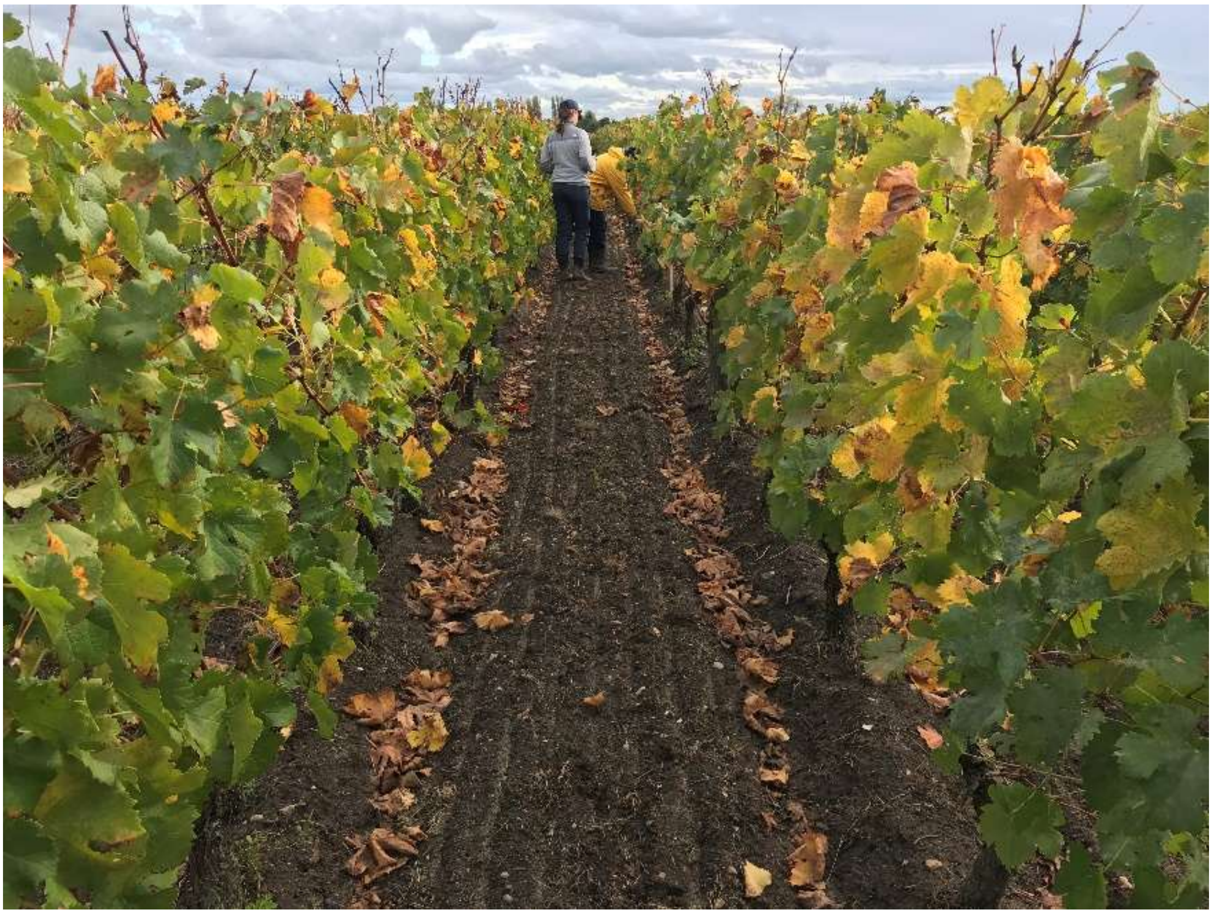

**Figure S10.** Illustration of the accumulation of grapevine leaf litter at the foot of the grapevines in a vineyard in the fall.

**Box S2. DNA extraction and ddPCR repeatability assessment**

In order to evaluate the repeatability of our method, we conducted additional analyses of soil samples. First, we randomly selected 44 soil samples out of the 318 samples that we collected in the experimental vineyard. For each of these samples, we took three portions of 2 g from the electric mixer bowl after the homogenization step. DNA was extracted from each portion separately, then the three portions were analyzed via ddPCR separately but at the same time (same operator, same day, same plate, and same machine run). The three portions of a sample were considered as extraction replicates (keeping in mind that the subsampling step's repeatability depends on the quality of sample homogenization). Next, we randomly selected another 42 soil samples from our collection of 318 samples. After a single DNA extraction per sample, we repeated the ddPCR analysis for three 4 µl aliquots of the DNA extraction product (previously diluted to the tenth) for each sample. The three aliquots of a sample were considered as ddPCR replicates. For both DNA extraction and ddPCR replicate data, we fit an analysis of variance model for the number of ITS copies as a function of the sample. In each case, we compared the residual variance (within-group variance, i.e. here the variance between replicates of a sample) to the variance explained by the model (between-groups variance, i.e. here the variance between samples) using an F-test, to evaluate the ability of our method to provide repeatable measures. In addition, we computed the coefficient of variation of the measurements associated with either DNA extraction or ddPCR as the ratio between the within-group standard-deviation (root square of the residual variance; expressed in the same unit as the response variable) and the mean number of ITS copies. The coefficient of variation indicates the mean dispersal of the values obtained for the replicates of a sample compared to the mean of all values, and a method is considered to have good repeatability when the coefficient of variation is inferior to 10%.

From the set of 44 samples randomly selected for DNA extraction repeatability assessment (Figure Box S2), we found 12.23 ITS copies per µl on average with a within-sample standard deviation of 4.60 ITS copies per µl. The analysis of variance indicated that the residual variance (within-sample variance SWS=21.12; associated to measurement error) was significantly lower than the explained variance (between-sample variance SBS=159.26 ; F=7.54; p=1.40 10-16) meaning that the error of measurement associated with our DNA extraction method was low compared to the between-sample variability. The within-sample coefficient of variation (*i.e.* associated with the repeated measurements of a sample) was equal to 41.2% on average, and varied from 7% to 160% with a median value at 30%.

From the 42 samples that were selected for the assessment of ddPCR repeatability (Figure Box S2), we obtained 11.83 ITS copies per µl on average, with a within-sample standard deviation of 2.68 ITS copies per µl. The analysis of variance indicated that the residual variance (SWS=7.17) was significantly low compared to the explained variance (SBS=334.01; F=46.59; p=1.09 10-40). The within-sample coefficient of variation (*i.e.* associated with the repeated measurements of a sample) was equal to 26.5% on average, and varied from 1% to 173% with a median value at 17%.

### Box S2 (continued). DNA extraction and ddPCR repeatability assessment

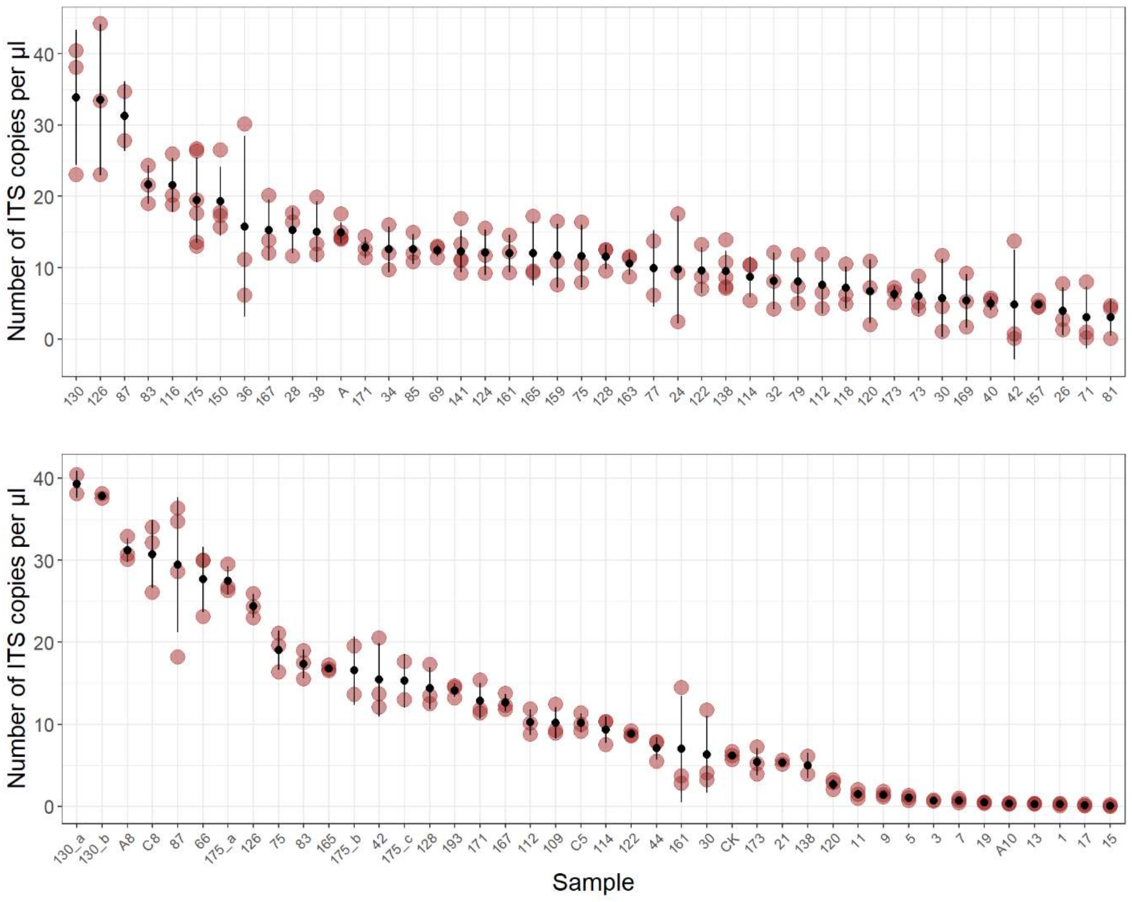


**Figure Box S2.** Triplicates of DNA extraction for 44 randomly selected soil samples (upper panel). Triplicates of ddPCR amplification for 42 randomly selected DNA samples (lower panel). Red dots correspond to repeated assessments of the number of ITS copies per µl found in the samples. Black dots and bars correspond to within-sample mean and standard deviation of the number of ITS copies per µl, respectively.

**Box S3. Comparison of primary inoculum spatial pattern with soil moisture and grapevine vigor spatial patterns**

Understanding the drivers of spatio-temporal variations in primary inoculum abundance is key to identifying levers for the preventive management of epidemic risk. At the leaf level, disease severity was found to be proportional to the amount of oospores formed per unit of leaf biomass (Maddalena *et al*. 2021). At the field scale, we hypothesized that the spatial variation in foliage disease severity explains the spatial variation in the abundance of primary inoculum because leaves (the oospore reservoir) remain close to the vine stock they fell from (*i.e.* a local accumulation of primary inoculum). Consequently, any spatially variable factor influencing downy mildew incidence over the course of the growing season could indirectly influence the distribution of primary inoculum in the soil. As an oomycete, *Plasmopara viticola* is particularly dependent on humidity at all stages of its life cycle (Gessler *et al*. 2011). For instance, soil moisture is required to trigger oospore germination and primary infections (Rossi and Caffi 2007), while a longer leaf wetness duration promotes leaf infections by downy mildew. Excessive grapevine vigor (defined as a propensity to assimilate, store, and/or use nonstructural carbohydrates for producing large canopies, associated with intense metabolism and fast shoot growth ; Hugalde *et al*., 2019) can result in increased susceptibility to disease, as high foliage density limits air movement and light penetration, as well as pesticide cover. In a study conducted in Germany, it was shown that canopy architecture can affect canopy microclimate and *in fine* susceptibility to grapevine downy mildew. More precisely, minimally pruned Chardonnay grapevines with large-volume canopies (*i.e.* vigorous) exhibited increased leaf wetness, higher relative humidity, and greater rates of downy mildew infection, as compared to intensively pruned grapevines with less dense foliage (Pennington 2019). Hence, spatial variation in soil moisture and grapevine vigor, for instance, could induce spatial variation in disease severity, resulting in spatial variation in soil primary inoculum concentration.

A private company (GEOCARTA) assessed soil electrical resistivity (SER) in the experimental vineyard in May 2006 using an ARP© system (see Andrenelli *et al*., 2013 for a detailed description of the device) which provided one measurement every 0.2 m along transects parallel to the rows. One measurement corresponded to the average of the electrical resistivity from the surface to 50 cm depth, *i.e.* apparent electrical resistivity. The raw data were first processed by 1D median moving window filtering along the acquisition profiles to eliminate noise in the measurements. Then, the dataset was interpolated in two dimensions on a 2 m-square grid (2D bicubic spline interpolation) by the company. We proceeded to a second, finer-scale interpolation so that final SER data corresponded to predicted values on a 0.1 m-grid. Soil electrical resistivity (the opposite of electrical conductance) is expected to decrease with increasing water content, as the charged ions in water make it highly electrically conductive. Therefore, we considered SER to reflect the inverse of soil moisture and we hypothesized that the distribution of primary inoculum was negatively related to the distribution of SER (and positively to soil moisture). We assumed that soil physico-chemical properties that control SER did not change significantly between the SER (2006) and inoculum measures (2022).

An optical captor (Trimble® GreenSeeker® rt100 crop sensing system, with AgGPS® FmX® acquisition system) was used to assess the normalized difference vegetation index (NDVI) of the foliage in the experimental vineyard during the growing season preceding soil sampling (*i.e.* June 2021). The optical captor was installed on a tractor and recorded one

**Box S3 (continued). Comparison of primary inoculum spatial pattern with soil moisture and grapevine vigor spatial patterns**

measure every 0.2 m along each row (see Goutouly *et al*., 2006 for a detailed description). NDVI is a widely accepted indicator of vegetative biomass and plant vigor, which we surmised to be related to modifications of microclimatic conditions favorable to downy mildew.

We investigated the relationship between spatial patterns in DM primary inoculum, SER and NDVI using cross variograms. First, we calculated a unique SER and a unique NDVI value for each primary inoculum sampling point of the regular grid as the mean of SER values within a 0.95 x 0.8 m rectangle around each sampling point (0.95 m being the distance between two vine stocks in a row, and 0.8 m being half the inter-row distance), and the mean of NDVI values within 0.48 m of each sampling point (0.48 m being half the distance between two vine stocks in a row), respectively. We reduced the skewness of the distribution of both SER and NDVI calculated data by applying a box-cox power transformation to each variable (Box and Cox 1964; Zhang *et al*. 1998). The interdependence between primary inoculum and SER or NDVI was examined using experimental cross-variograms, estimated by the following:

$$\hat{\gamma}_{uv}(\mathbf{h}) = \frac{1}{2m(\mathbf{h})}\sum_{i=1}^{m(\mathbf{h})}[\{z_u(\mathbf{x}_i) + z_u(\mathbf{x}_i + \mathbf{h})\} \{z_v(\mathbf{x}_i) + z_v(\mathbf{x}_i + \mathbf{h})\}].$$

where $m(h)$ is the number of comparisons at lag $h$; $z_u(x_i)$ and $z_u(x_i + h)$ are the measured values of the variable $z_u$; $z_v(x_i)$ and $z_v(x_i + h)$ are the measured values of $z_v$ at $x_i$ and $x_i + h$, respectively. A positive relationship between the cross semivariance and distance lag indicated a positive coregionalization between both variables.

The cross-variograms revealed a clear negative coregionalization between downy mildew primary inoculum and SER (Figure 1), or alternatively a positive coregionalization with soil moisture. This result is consistent with our hypothesis that soil moisture promotes disease incidence locally, which in turn explains local inoculum accumulation.

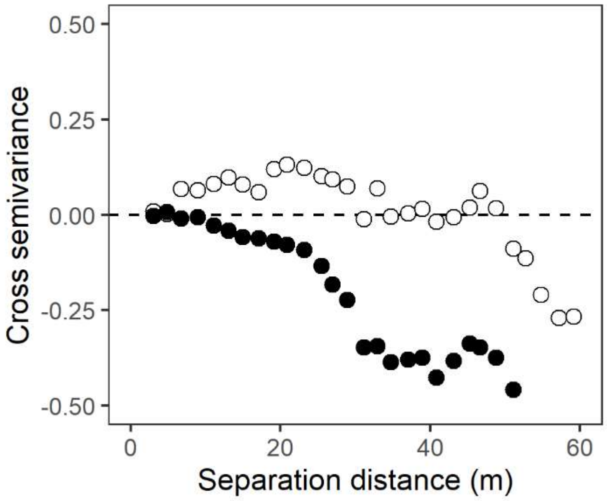


**Figure Box S3**. Cross-variograms of downy mildew primary inoculum vs soil electrical resistivity (black) or grapevine NDVI (white).

**Box S3 (continued). Comparison of primary inoculum spatial pattern with soil moisture and grapevine vigor spatial patterns**

The cross-variogram of downy mildew primary inoculum *vs* grapevine vigor did not indicate a clear coregionalization of the two variables (Figure 1; weak positive correlation up to 25 m of distance then weak negative correlation). This result did not support our hypothesis for a cascading effect of vigor on local disease incidence (due to greater susceptibility).

We demonstrated that this distribution pattern could be indirectly determined by the spatial variability of some abiotic factors that control disease incidence. Indeed, we showed that the concentration of primary inoculum in the soil was positively coregionalized with soil moisture. It has been reported that soil moisture was also one of the main factors explaining soilborne disease occurrence (Jiang *et al*. 2021). These findings support the hypothesis of local accumulation of inoculum, *i.e.* the amount of inoculum formed in a given area in a field is proportional to disease incidence, then leaves fall and remain close to the vine they fell from and oospores are unlikely to disperse once in the soil matrix. As mentioned above, this local accumulation pattern can be reinforced by soil ridging. The absence of a clear relationship with grapevine vigor may be due to several reasons. Contrary to soil physical characteristics, grapevine vigor is dependent on many factors and may vary rapidly during the course of the growing season. Therefore, it is more difficult to establish a relationship between grapevine vigor and accumulated inoculum in the soil, especially as we tried to relate a discrete assessment of vigor with an assessment of downy mildew primary inoculum that reflects disease severity integrated over the whole season. Second, the detection of a relationship between grapevine vigor and primary inoculum concentration may have been mitigated by the phytosanitary treatments applied in the experimental field during the 2021 growing season. Therefore, grapevine vigor or foliage density cannot be used to predict DM primary inoculum distribution, and controlling for vigor is not a promising lever for DM primary inoculum management unless further research establishes a stronger relationship.

**Box S3 (continued). Comparison of primary inoculum spatial pattern with soil moisture and grapevine vigor spatial patterns**

**References (continued)**

Jiang, G., Wang, N., Zhang, Y., Wang, Z., Zhang, Y., Yu, J., et al. 2021. The relative importance of soil moisture in predicting bacterial wilt disease occurrence. Soil Ecol. Lett. 3:356–366.

Maddalena, G., Russo, G., and Toffolatti, S. L. 2021. The study of the germination dynamics of *Plasmopara viticola* oospores highlights the presence of phenotypic synchrony with the host. Front. Microbiol. 12:698586.

Pennington, T. 2019. Natural pest suppression in vineyards under innovative management. PhD thesis.

Rossi, V. and Caffi, 2007. Effect of water on germination of *Plasmopara viticola* oospores. Plant Pathology, 56, 957-966.

Zhang, C., Selinus, O., and Schedin, J. 1998. Statistical analyses for heavy metal contents in till and root samples in an area of southeastern Sweden. The Science of The Total Environment. 212:217–232.
